# Changes within neural population codes can be inferred from psychophysical threshold studies

**DOI:** 10.1101/2020.03.26.010900

**Authors:** Jason Hays, Fabian A. Soto

## Abstract

The use of population encoding models has come to dominate the study of human visual neuroscience, serving as a primary tool for making inferences about neural code changes based on indirect measurements. A popular approach in computational neuroimaging is to use such models to obtain estimates of neural population responses via inverted encoding modeling. Recent research suggests that this approach may be prone to identifiability problems, with multiple mechanisms of encoding change producing similar changes in the estimated population responses. Psychophysical data might be able to provide additional constraints to infer the encoding change mechanism underlying some behavior of interest. However, computational work aimed at determining to what extent different mechanisms can be differentiated using psychophysics is lacking. Here, we used simulation to explore exactly which of a number of changes in neural population codes could be differentiated from observed changes in psychophysical thresholds. Eight mechanisms of encoding change were under study, chosen because they have been proposed in the previous literature as mechanisms for improved task performance (e.g., due to attention or learning): specific and nonspecific gain, specific and nonspecific tuning, specific suppression, specific suppression plus gain, and inward and outward tuning shifts. We simulated psychophysical thresholds as a function of both external noise (TvN curves) or stimulus value (TvS curves) for a number of variations of each one of the models. With the exception of specific gain and specific tuning, all studied mechanisms produced qualitatively different patterns of change in the TvN and TvS curves, suggesting that psychophysical studies can be used as a complement to inverted encoding modeling, and provide strong constraints on inferences based on the latter. We use our results to provide recommendations for interested researchers and to re-interpret previous psychophysical data in terms of mechanisms of encoding change.

## Introduction

The use of population encoding models has come to dominate the study of human visual neuroscience, serving as a primary tool for making inferences about neural code changes based on indirect measurements, such as psychophysical [e.g., 1, 2, 3, 4, 5, 6, 7, 8] and neuroimaging measures [e.g., 9, 10, 11, 12, 13, 14, 15]. One of the primary benefits of these models is that they can be applied to understand the neurocomputational mechanisms of perceptual processes when more invasive methods are not easily available, as is the case in most human neuroscience studies. Population encoding models make a number of assumptions that constrain the space of possible inferences that one can make from limited data [13]. Researchers use these models to make inferences about how neural codes change from a baseline state during and after certain cognitive events [15].

The standard population encoding model (see Figure 1a, under the title “Encoder”, [16, 17, 18, 4]) consists of a population of neural channels (representing a neuron or a population of neurons with similar selectivity), each characterized by a tuning function that responds more strongly to stimuli that have features similar to its preferred stimulus (i.e., a specific orientation). When a stimulus is presented to the encoder, the set of channels outputs a population response (i.e., a vector of response rates; see Figure 1a), which is affected by internal noise (error bars in the figure). Information about the stimulus is distributed across neural channels in the population response, so that when it is needed for a behavioral task, a decoder must integrate it and recover it (see Figure 1a, Decoded Stimulus).

**Figure 1:**
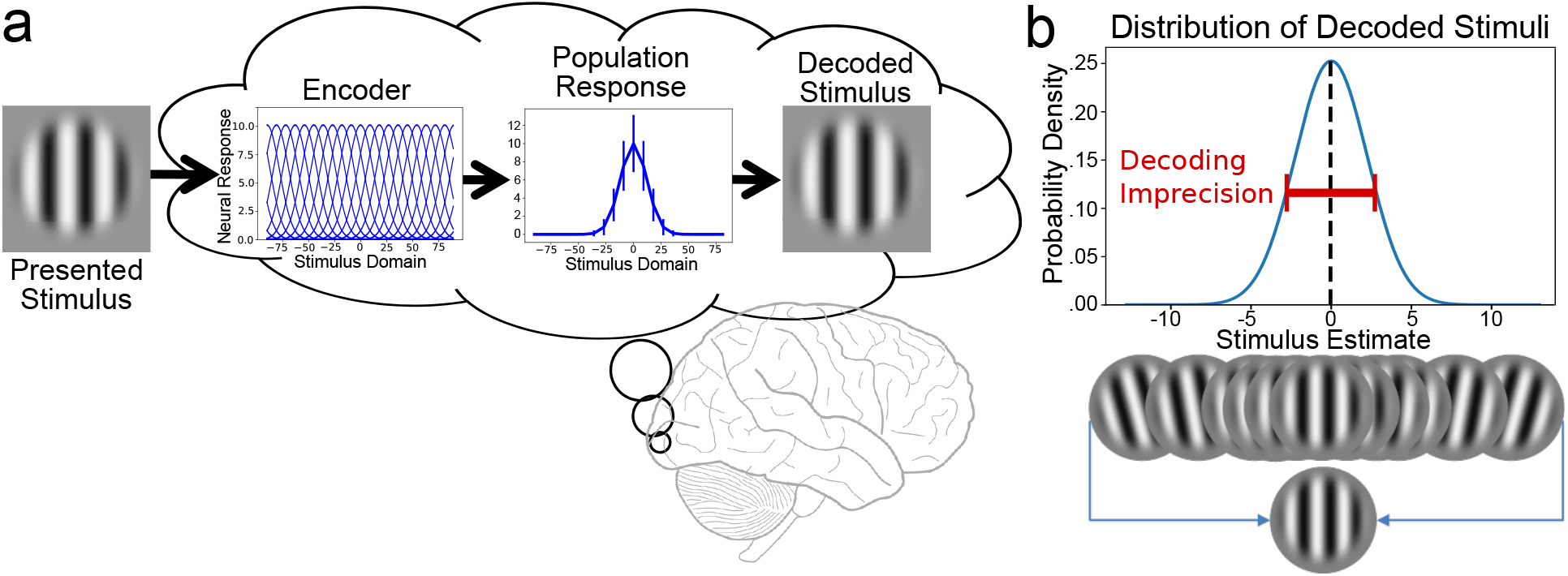
(a) Encoding and decoding of grating orientation. The target stimulus is encoded by a population of neural channels to produce a noisy population response (the error is reported in standard deviations), which is then decoded to produce a single, noisy stimulus estimate. (b) Using optimal decoding, stimulus estimates from many presentations of the same stimulus will be normally distributed around the true stimulus value. The width of the distribution represents decoding imprecision.

An important problem is exactly how the brain decodes stimulus information from distributed population responses, when such information is needed for a behavioral task. There are many possible decoding schemes [e.g., 19, 20, 21, 22], and some choices lead to an inherent ambiguity in whether a behavioral observation is due to encoding changes versus decoding changes [23]. Confronted with this dilemma, many authors assume optimal decoding via maximum likelihood estimation (MLE; e.g., [1, 24, 3, 4, 7, 6]). There is usually a single optimal solution for a well-posed statistical problem, and thus this solves the issue of ambiguity regarding whether a change in behavior is due to changes in encoding versus decoding processes. There are two additional advantages of using an optimal decoder. First, its use has proven to be fruitful in the previous literature, allowing researchers to make links between psychophysical measures and known properties of neural encoding [e.g., 6]. Second, the choice of an optimal decoder facilitates linking neural encoding mechanisms to psychophysical measures, particularly when signal detection theory is used as a measurement framework. More specifically, because neural noise influences the population response shown in Figure 1a, the decoded stimulus estimate is also noisy, following a particular probability distribution, as shown in Figure 1b. When using optimal decoding through MLE, this is a normal distribution centered at the true stimulus value and having a standard deviation, or width, that represents decoding imprecision (see Figure 1b).

The properties of the encoding channels directly affect the population response to a particular stimulus, which in turn affects decoding imprecision; thus, decoding imprecision is linked to the attributes of the encoding channels (see Figure 2). Changes to neural codes produce concomitant changes in the precision of decoding, and therefore the discriminability of stimuli from the point of view of signal detection theory. Depending on how the neural codes are altered, they can worsen or improve the discriminability of stimuli differently across the stimulus domain. For example, increasing the responsiveness of channels at the target usually improves the discriminability around the target and for nearby stimuli. On the other hand, decreasing the width of channels at the target may improve the discriminability around the target, but decrease discriminability for nearby stimuli. Because different neural changes produce different outcomes, our goal here was to determine through simulation whether the different mechanisms of change proposed in the neuroscientific literature could be distinguished using psychophysics. While previous researchers have distinguished between a couple of mechanisms [3], nobody has attempted to distinguish among all of those often proposed in the literature.

**Figure 2:**
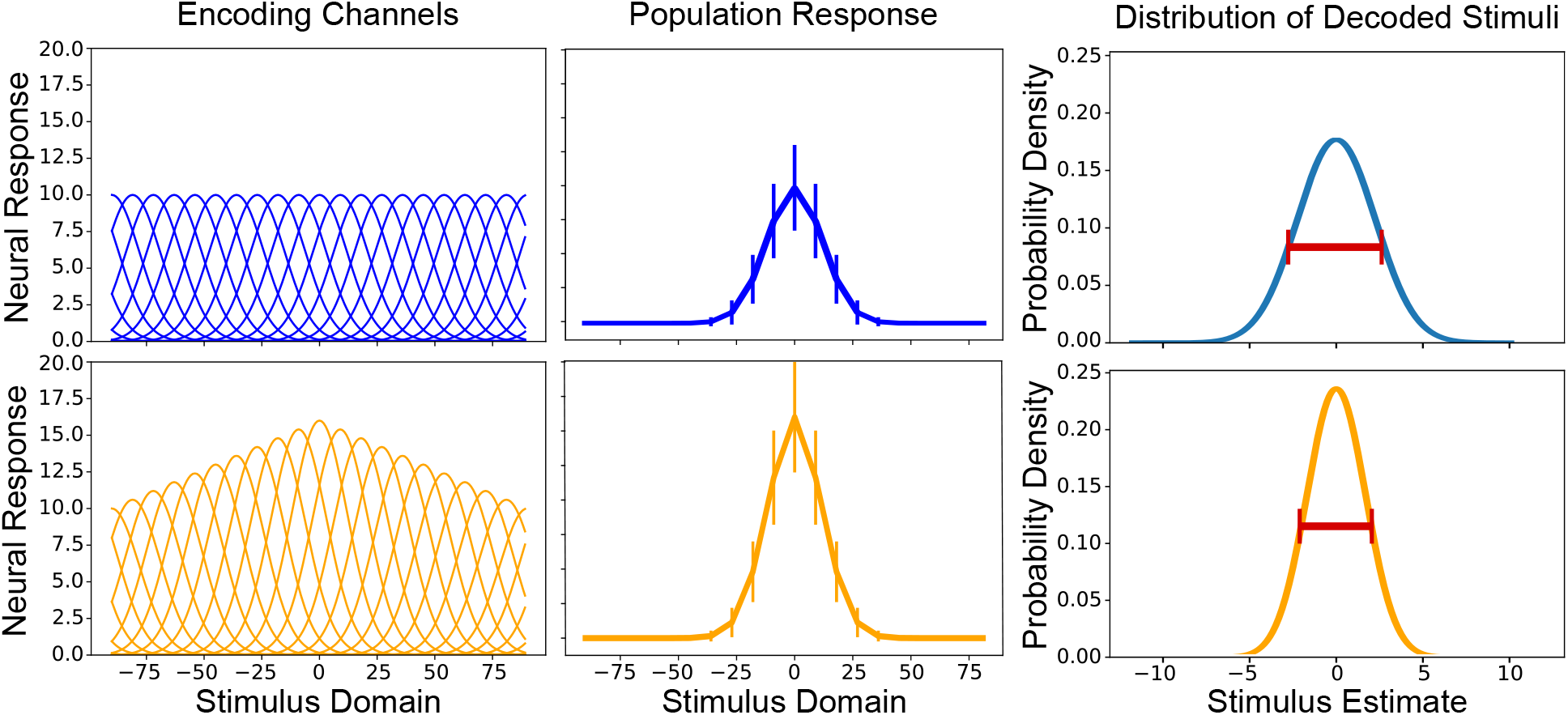
Relation between properties of neural encoding and decoding imprecision. Columns represent 1) the encoding channels, 2) the resulting population response, and 3) the final distribution of decoded stimuli, when the target stimulus presented is at zero. Compared to a baseline homogeneous condition (top, in blue), an increase in responsiveness of neural channels around the target stimulus (bottom, in orange) leads to a stronger population response and to more precise decoding (as measured by the red bar representing full-width at half-maximum of the distribution of stimulus estimates).

As indicated earlier, there have been two main ways in which researchers have applied the encoding/decoding model presented in Figure 1. First, they have used the encoding model to infer properties of neural representations from neuroimaging data. For example, inverted encoding modeling [10, 15, 13] assume a homogeneous population encoding model, like that presented in blue in the top-left panel of Figure 2. A range of stimuli are presented to the model to obtain average population responses, like those presented in the middle panel of Figure 2. Such responses are then used as predictors of neuroimaging data using multivariate linear regression and, after inversion of the model, new datasets are used to recover estimates of population responses under multiple conditions [15]. For example, a model trained with neuroimaging data during a no-task baseline (i.e., only stimuli are presented) could be used to recover and compare population responses from the no-task and a task condition (e.g., stimuli must be attended, categorized, etc.).

Despite its successes, inverted encoding modeling makes rather strong implicit assumptions about encoding (e.g., homogeneous population codes, normal neural noise) and about the link between neural activity and neuroimaging measures (a linear measurement model with additive normal noise at each measurement, and independent across measurements) [see 14, 8]. In addition, the approach has known identifiability problems. Some researchers [12] have pointed out that inverted encoding modeling sometimes cannot tell apart different changes at the level of encoding channels (see Figure 2, left panel), which produce similar changes at the level of recovered population responses. Others [15] have pointed out that the approach is aimed at studying the level of population responses (see Figure 2, middle panel), and not the level of underlying mechanisms of encoding change. However, one may be interested in selecting among a limited number of possible mechanisms of encoding change, and reach conclusions that can be linked to work using direct measures of neural activity. In that case, the goal would be to use recovered population responses to make inferences about the underlying code.

A second way in which the encoding/decoding model can be used to study human vision is by inferring properties of neural representations from psychophysical data. As indicated above, changes in neural encoding produce concomitant changes in the precision of neural decoding. Psychophysical thresholds can provide estimates of such precision, and obtaining thresholds as a function of some stimulus property or condition might provide strong constraints on mechanisms of encoding change. For example, **(author?)** [3] obtained psychophysical thresholds for the detection of motion direction across levels of external noise, to distinguish whether changes in height or tuning of population responses underlie selective attention. Their application of the encoder/decoder model is particularly interesting, as their goal was essentially the same as the stated goal of inverted encoding modeling [15].

Unfortunately, many other psychophysical studies have been much less clear about their inferential goals. A common problem is that several techniques have been developed to estimate psychophysical functions that look similar to either neural channel tuning functions or population responses. Indeed, some researchers highlight analogous findings in the psychophysical and neurophysiological literatures, directly comparing properties (e.g., width) of behavioral tuning functions against similar properties of tuning functions obtained from single-neuron electrophysiology [e.g., 25, 26, 27].

To summarize, a mechanism of encoding change produces changes at the three levels shown in Figure 2: at the level of neural channel tuning functions (left panel in the figure), at the level of the population response to a stimulus (middle panel in the figure), and at the level of decoding precision measured through psychophysics (right panel in the figure). Until recently, most researchers in both the neuroimaging and psychophysical literatures have been unclear about exactly what level is the target of their inferences, although in some exceptional cases researchers make clear that this is the level of population responses [3, 15]. An open question is whether different mechanisms of encoding change (i.e., at the level of neural channels) that have been highlighted in the literature may produce different patterns of results at the level of population responses and/or decoding precision (i.e., psychophysical thresholds).

In the present project, we were particularly interested in the less explored area of psychophysics, as knowledge about exactly what inferences can be made about mechanisms of encoding change from such data is useful in two ways. First, because psychophysical data may be used as a supplement to inverted encoding modeling of neuroimaging data, helping to narrow down the range of possible models underlying a particular behavior. This is specially true when multiple mechanisms can give rise to similar patterns in the neuroimaging data [12]. Second, because such knowledge could help in the interpretation of psychophysical results in terms of changes in neural encoding. As indicated earlier, although it is common for researchers to draw links between psychophysics and neural codes, in many cases this is not done using a formal framework. One of our goals was to provide general guidelines to choose psychophysical design and interpret their results in terms of mechanisms of encoding change. Such guidelines could also apply to the re-interpretation of previous psychophysical results, from studies that were either not focused on encoding mechanisms [e.g., 28, 29], limited the number of mechanisms of encoding change explored [e.g., 3], or did not use a formal framework to link psychophysical functions with properties of neural codes [e.g., 25, 27].

We used simulation work to explore exactly which of a number of changes in neural population codes could be differentiated from observed changes in psychophysical thresholds. We focus on the simple encoding/decoding model presented in Figure 1, which has been the focus of much previous research. We simulate a variety of encoding changes from a homogeneous population baseline, while assuming an optimal decoder. We focus on whether changes in that baseline can be detected through a variety of psychophysical measures.

Figure 3 provides one example for each of the mechanisms of encoding change that were explored in the current simulation work. In this and all other figures, we refer to the main stimulus of interest as the target (i.e., we assume that it is specifically targeted by some experimental manipulation, such as training or cueing), which in all cases has a value of zero. In each simulation, the baseline encoding population (i.e., the blue homogeneous population in the top-left panel of Figure 2) was modified by a different mechanism of encoding change. Each mechanism affects one of the three parameters of the tuning function (width, height, and center or position; see Figure 4) differently. While specific gain, nonspecific gain, and specific suppression all affect the height of tuning functions, specific gain boosts the height for channels with position nearest to the target more than others, specific suppression decreases the height of channels farthest from the target more than others, and nonspecific gain indiscriminately boosts the height of all channels. Specific tuning and nonspecific tuning narrow the width of the tuning functions, but specific tuning narrows the width of channels nearest to the target more than others, while nonspecific tuning indiscriminately narrows the width of all channels. Inward and outward tuning shifts affect the position parameters of the tuning functions; they are “specific” mechanisms insofar as they move each channel’s position depending on its distance from the target. Specific suppression plus nonspecific gain implements both mechanisms simultaneously.

**Figure 3:**
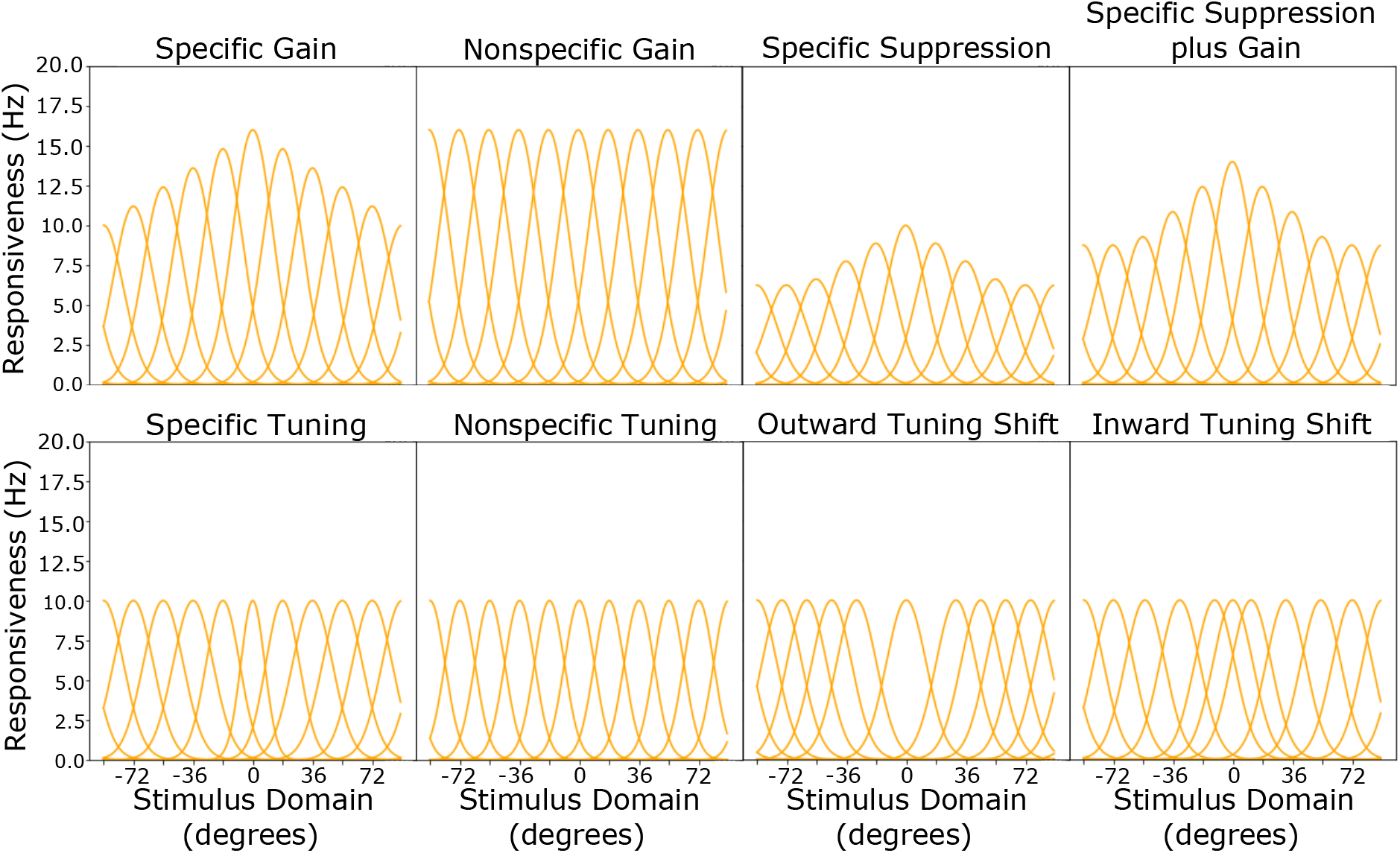
In addition to the homogeneous baseline depicted in the top-left panel of Figure 2, there were eight different mechanisms of encoding change from baseline under study. Each mechanism is expected to produce unique patterns of thresholds in psychophysical experiments.

**Figure 4:**
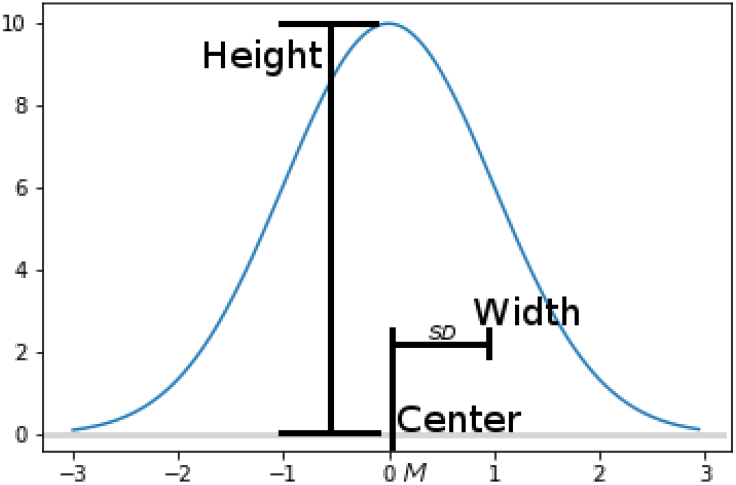
Gaussian curves are used to describe the shape of channel tuning functions. These are defined by three parameters: width (standard deviation), height (maximum average responsiveness, in Hz), and center or position (mean).

These mechanisms of encoding change were selected because they have been proposed in the previous literature to improve task performance in at least some circumstances. Both gain and tuning have been discussed in the attention, conditioning, and category learning literatures [25, 30, 9, 31, 32, 33, 34, 35, 36]. Specific suppression and specific suppression plus gain have been proposed as external noise reduction mechanisms of attention [37, 26]. Outward tuning shifts have been suggested as mechanisms for improving the performance around category bounds during category learning [38]. Inward tuning shifts have been suggested as a mechanism of associative learning [39, 40] that causes conditioned stimuli to be over-represented.

## Results

In this study, we used an encoding-decoding observer model (see Figure 1) to simulate how a number of mechanisms of encoding change (see Figure 3) would affect decoding imprecision, as measured through sensitivity thresholds. We used Gaussian tuning functions to represent the selectivity of encoding channels to different stimuli, independent Poisson neural channel noise, and statistically optimal decoding through maximum likelihood estimation (MLE). The Gaussian tuning functions had three parameters (width, height, and center or position; see Figure 4), which were each influenced differently by the mechanisms of encoding change. The width is implemented as the standard deviation of the Gaussian, the height is the average responsiveness at the preferred stimulus, and the position represents the preferred stimulus, implemented as the mean of the Gaussian. We created a baseline, homogeneous population encoding model, which had tuning functions with a width of 12 degrees, a height of 10 Hz, and positions evenly spread along a circular stimulus domain [−90,90). We simulated both a sparse population with 10 channels and a dense population with 20 channels. Neural channel noise was simulated using independent Poisson noise, a common choice in the previous literature [e.g., 1, 3, 4]. Statistically optimal decoding was implemented via MLE, which produces a normal distribution of stimulus estimates, centered at the true stimulus value and with a standard deviation proportional to the sensitivity threshold (see Figure 1b; for more details, see *The encoding-decoding observer model* in the *Methods* section).

Starting from the homogeneous baseline, we implemented eight different mechanisms of encoding change, which include nonspecific gain, nonspecific tuning, specific gain, specific tuning, inward tuning shift, outward tuning shift, specific suppression, and specific suppression plus gain (the only combination of mechanisms). The effect of each one of these mechanisms on the tuning functions of the population of channels is exemplified in Figure (3). We ran a total of 244 simulations, each characterized by a different population encoding model. Monte Carlo simulations were used to estimate each threshold, with 50,000 iterations used per threshold. In one set of simulations, we manipulated the external noise (0 to 0.5) to influence the thresholds, and in the other set of simulations, we manipulated the presented stimulus (−90 to 90) to affect the thresholds. For more details on the implementation of each model, the Monte Carlo estimation, and experimental manipulations, see the *Methods* section.

For each set of simulations, we assessed under what conditions the simulated thresholds fell above or below those of the baseline. We expected each neural change to create a qualitatively different pattern of thresholds in our simulated experimental paradigms (explained below). We assumed that increasing or decreasing the magnitude of any given neural change would only affect the magnitude of the differences with baseline.

### Population Response Curves

As indicated earlier, one goal of our work was to encourage researchers that use inverted encoding modeling to supplement their studies with psychophysical data, as a way to narrow down the the range of possible models underlying a particular behavior of interest. To frame our work in those terms, we started by checking to what extent the different mechanisms of encoding change under study could be differentiated via inverted encoding modeling.

As indicated above, the main goal of inverted encoding modeling is to obtain estimates of population responses [15], such as those shown in the middle panels of Figure 2. For this reason, we started by obtaining such population responses after presentation of the target stimulus, from each one of the models under study. The obtained responses are shown in Figure 5. Note that these are the true mean population responses without any noise, and thus what inverted encoding models aim to estimate.

**Figure 5:**
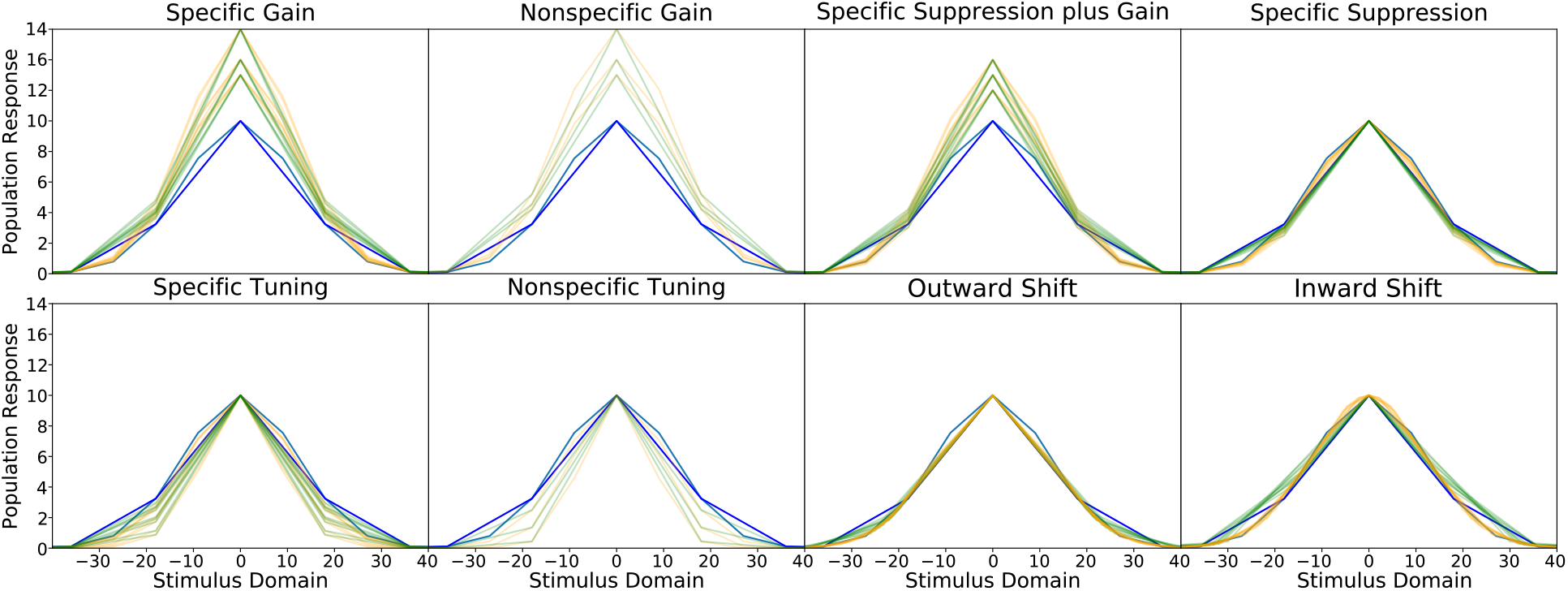
Population responses obtained from each of the encoding models under study. Blue curves correspond to baseline models, orange curves to encoding changes implemented in a dense population (20 channels), and green curves to encoding changes implemented in a sparse population (10 channels).

Each one of the population responses shown in Figure 5 were fitted to a model of the population response curve (see Equation 8), which has the Gaussian function as a special case, but that can also produce curves with lower or higher curvature at the peak, including exponential functions (without any curvature at the peak). The fitted curve is similar to that used in previous inverted encoding modeling [e.g., 38], in that it allows to obtain separate estimates of the population response’s height and width, but also provides estimates of the curvature (i.e. width) of the curve at the peak. The fit of the population response curve was in most cases excellent. Figure 6 shows that fit for responses obtained from sparse models. The inserts at the top-left of each plot show the distributions of *r*^2^ values obtained for that model, with dots representing individual values, and the diamond representing the mean value. The main plot shows an example comparison between a fine-detail population response and the prediction from the population curve fitted to sparse data, for the model closest to the mean.

**Figure 6:**
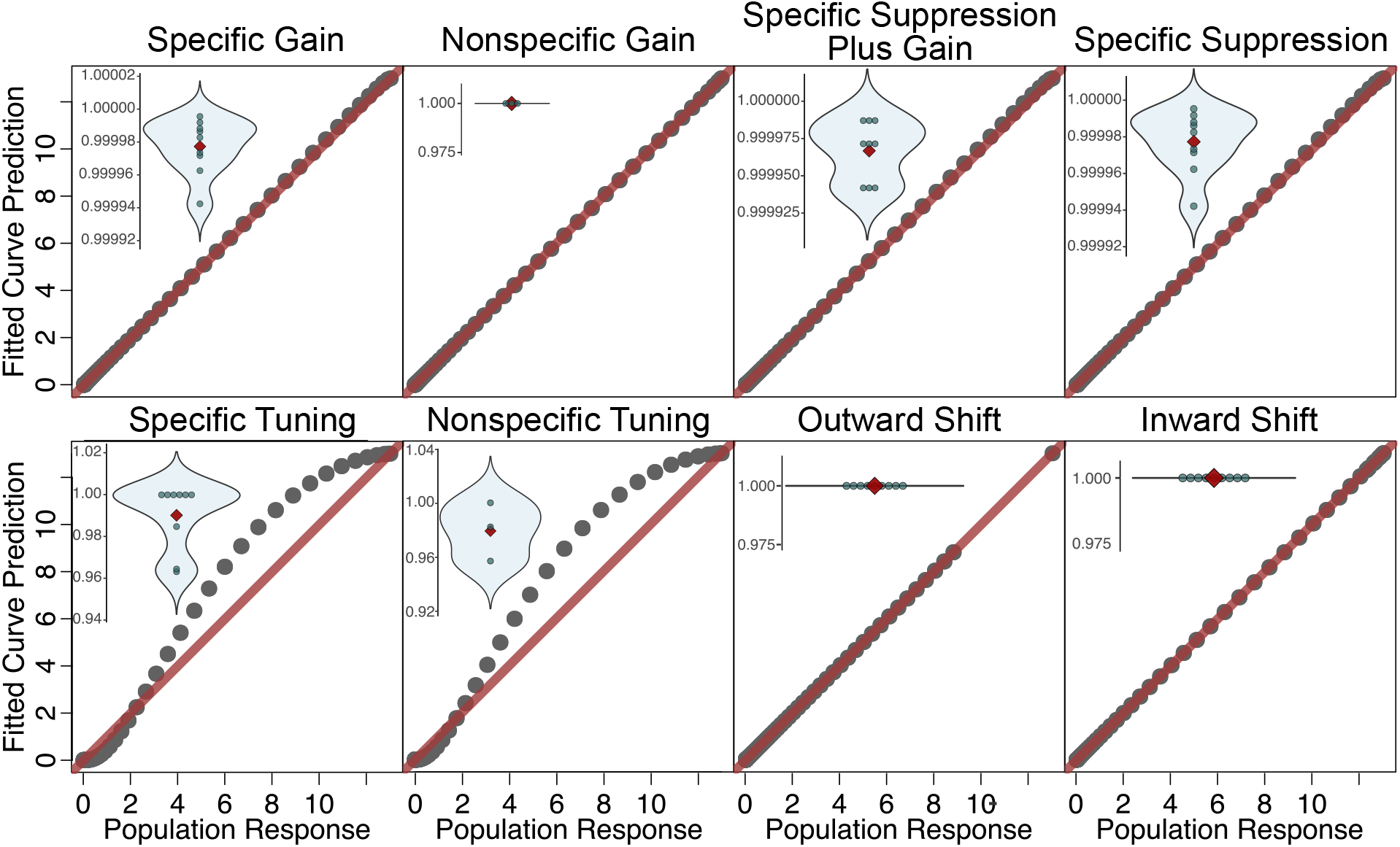
Fit of the population response curve to the mean population responses obtained from the models. The main plots compare the smooth population response obtained from a model with high density of channels (see Figure 8) in the *x*-axis, against the predicted values for the same population response obtained from a curve fitted to a sparse population response, with low density of channels. The inserts at the top-left of each plot show the distributions of *r*^2^ values obtained for that model, with dots representing individual values, and the diamond representing the mean value. Axes of the main plots have been re-scaled to ease comparison across models.

First, note that most of the *r*^2^ values are higher than 0.99, indicating excellent fits. Only specific tuning and nonspecific tuning provide relatively lower values, which never drop below an *r*^2^ of 0.96. The values obtained from dense population models (20 channels) were all so high that were rounded up to 1.00 due to machine precision. This indicates that, in most cases, our model of the population curve provided a good description of population response obtained from the models and we can interpret the shape of population responses through the recovered parameters.

Such recovered parameters were subtracted from the corresponding parameters obtained from baseline models, and the distribution of such differences are shown in Figure 7. We must first note that many applications of inverted encoding modeling focus simply on whether the population response changes in height versus width in a given experimental condition (e.g., attention). It can be seen from the Figure that while an increase in height of the curve was always diagnostic of a gain mechanism at the neural encoding level (specific gain, nonspecific gain, or specific suppression plus gain), changes in the width of a curve are not very diagnostic of changes at the neural encoding level. We see narrowing of the curve width due both to changes in tuning (specific and nonspecific) and other mechanisms (specific gain, specific suppression plus gain, specific suppression). This mirrors the point made by previous simulation work [12] showing that changes in tuning of recovered population responses are not diagnostic of changes in tuning of encoding neural channels. A final result is that tuning shifts (i.e., changes in the preferred stimuli of individual channels) cannot be directly observed from population responses, which look identical to those observed during baseline. This observation is in line with published results, which showed that recovered population responses do not reveal an outward tuning shift, but that such a mechanism can be inferred by using special data analyses [38] (we return to this study in the discussion section).

**Figure 7:**
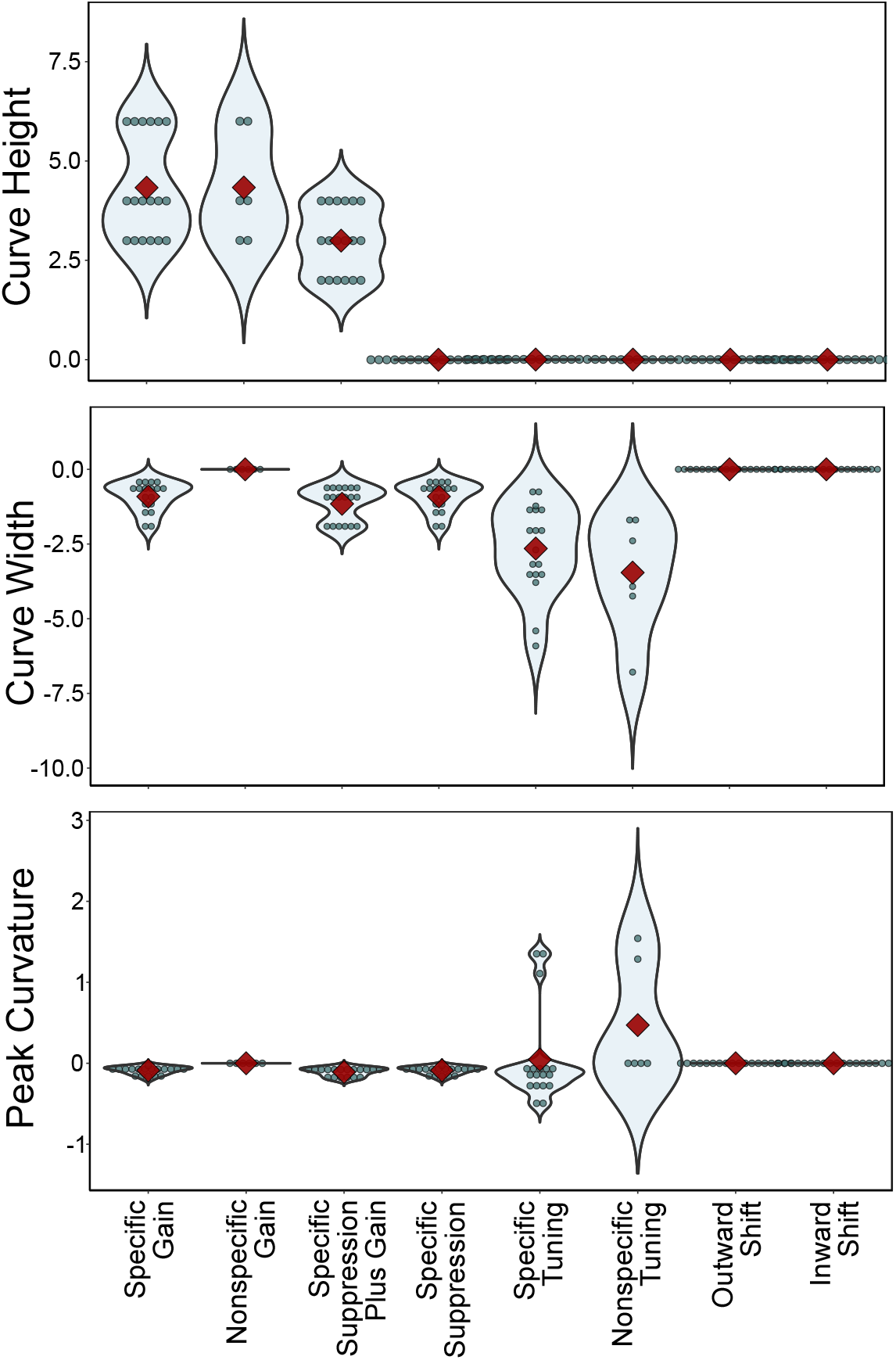
Distributions of parameters recovered from the population response to the target stimulus, presented separately for each model under study. The values represent the difference in estimated parameter between a particular model and its corresponding baseline, with values higher than zero representing “taller” or “wider” curves, and values below zero representing “shorter” or “narrower” curves. Each dot represents a different model variation and the red diamond represents the mean of the distribution.

What is most revealing about the results shown in Figure 7 is that they do reveal some information about the underlying mechanisms of encoding change, even though they provide that information only indirectly. For example, nonspecific gain produces in all cases a very specific pattern, with both width parameters identical to those seen at baseline, and the curve height parameter higher than baseline. However, most cases do seem difficult to tell apart from one another, which seems to be in line with the recommendation of not making inferences about underlying channel tuning functions from recovered population responses [15].

On the other hand, we would be amiss if we did not point out that the inverted encoding modeling approach has more potential to provide information about encoding changes than what current practice permits. First, note that the parameters provided in Figure 7 were obtained from population responses to a single target stimulus. In real applications, multiple stimuli with different values in the dimension of interest are presented, the population responses are estimated for each one of them, shifted to have a common zero mean, and averaged to obtain a single estimate of the population response. This practice allows to obtain better estimates of the population response *only when nonspecific mechanisms are involved.* To understand why this is the case, Figure 8 shows the population responses obtained by presenting seven evenly-spaced stimuli, starting from the target and moving towards the right side of the dimension, to an ultra-dense version of the population encoding models used here. This allowed to obtain smooth population responses showing all the information ideally available from an inverted encoding modeling experiment. Note first that when nonspecific gain or tuning are involved, all population responses have the same shape and their average is a good estimate of that curve. For all other cases, population responses vary with presented stimulus. For example, in specific gain population responses drop in height as the stimulus gets away from the target, in specific tuning population responses widen as the stimulus gets away from the target, and so on.

**Figure 8:**
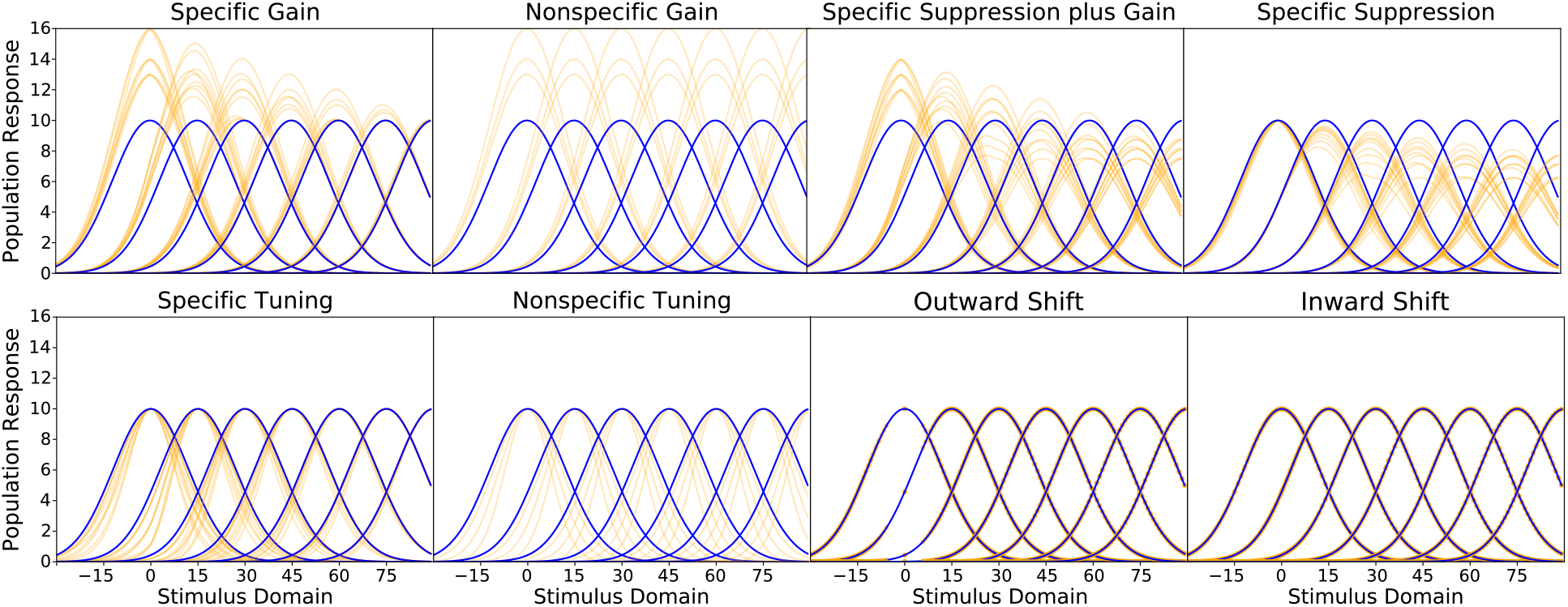
Idealized population responses obtained from ultra-dense population encoding models. They show that the current practice of averaging population responses from different stimuli reduces the information available about underlying mechanisms of encoding change.

Thus, the common practice of averaging shifted estimates of population responses has three undesirable consequences. First, when a specific mechanism is involved, averaging produces biased estimates of any of the true underlying population responses. Second, averaging may reduce the size of the effect of an experimental factor on population responses, and in some cases it might even artificially get rid of such effect (e.g., when averaging responses higher and lower from baseline, in the case of specific suppression plus gain). Finally, averaging discards an important amount of information about the underlying mechanism of encoding change. For example, the parameters recovered from a single curve in Figure 7 cannot distinguish between specific gain and specific suppression plus gain, but the latter mechanism is the only one that produces suppressed population responses away from the target in Figure 8. Similarly, specific suppression, specific tuning, and nonspecific tuning cannot be distinguished from the parameters shown in Figure 7, but they clearly produce different patterns of population responses as a function of stimulus value in Figure 8.

Thus, it seems like the inability to differentiate mechanisms of encoding change using inverted encoding modeling is in part due to the practice of averaging population responses obtained by presentation of different stimuli. A much better approach to differentiate between the different mechanisms in Figure 3 would be to estimate a different population response for a number of stimulus values along the dimension. This, however, would require many times the amount of data that is usually obtained in inverted encoding modeling studies, and might not solve other identifiability issues highlighted by recent research [12]. In addition, traditional inverted encoding analyses cannot provide information about tuning shift mechanisms, and it is not clear whether special analyses proposed to obtain such information for outward tuning shifts [38] would similarly work to infer inward tuning shifts.

Thus, we now turn to the main question of our current research, which is whether or not mechanisms of encoding change can be distinguished from one another based on psychophysical data alone.

### Threshold vs Noise (TvN) Curves

The equivalent-noise paradigm [41] is a widely-used psychophysical tool that involves measuring the sensitivity of the visual system to changes in a given variable (e.g., grating orientation) at many different values of external noise that is added to the stimulus. Sensitivity is usually measured via thresholds, which as indicated earlier are estimates of decoding imprecision. Figure 9 shows an explanation of the resulting Threshold versus external Noise (TvN) curve (also sometimes called TvC curve, where C stands for “Contrast Noise”). The typical shape of this TvN curve has a flat section at low levels of external noise, where performance is almost exclusively limited by internal sources of noise, such as neural noise in our simulations. This is followed by a curved section where external noise starts exerting its influence, and ends with a linearly-increasing section where performance is almost exclusively determined by the level of external noise.

**Figure 9:**
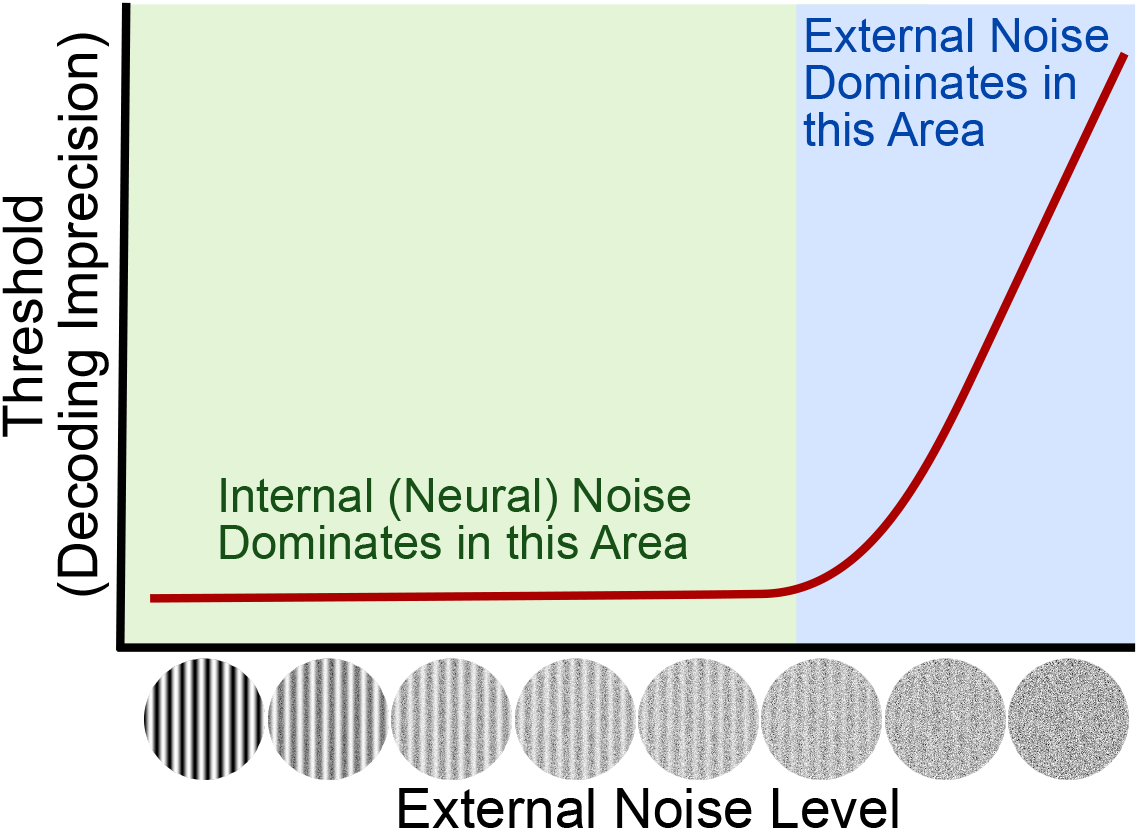
Explanation of a Threshold versus external Noise (TvN) curve.

TvN curves have been used in the past to characterize psychophysical observer models [42], as well as encoding/decoding observer models like the one presented in the introduction [e.g., 1, 3]. In the context of our study, the TvN curve has been shown to provide useful information about mechanisms of change in encoding populations. The reason for its usefulness is that different changes in encoding should produce different changes in the curve, depending on whether they affect internal or external noise. For example, **(author?)** [3] found through simulations that a nonspecific gain mechanism of attention would specifically suppress internal noise, making the TvN curve drop only at low levels of external noise (green area in Figure 9). On the other hand, a “tuning” mechanism of attention (what we have called specific suppression with nonspecific gain) would specifically suppress external noise, making the TvN curve drop only at high levels of external noise.

The goal of the present set of simulations was to expand previous work [1, 3] and determine how *all* the changes in population codes shown in Figure 3 affect the shape of the TvN function. The obtained TvN curves are shown in Figure 10, with each panel representing a different mechanism of encoding change. Baseline curves are shown in blue, whereas curves obtained from the models in Figure 3 are shown in either orange (dense population) or green (sparse population). In the rest of this section, we describe the results shown in Figure 10 in detail for each model.

**Figure 10:**
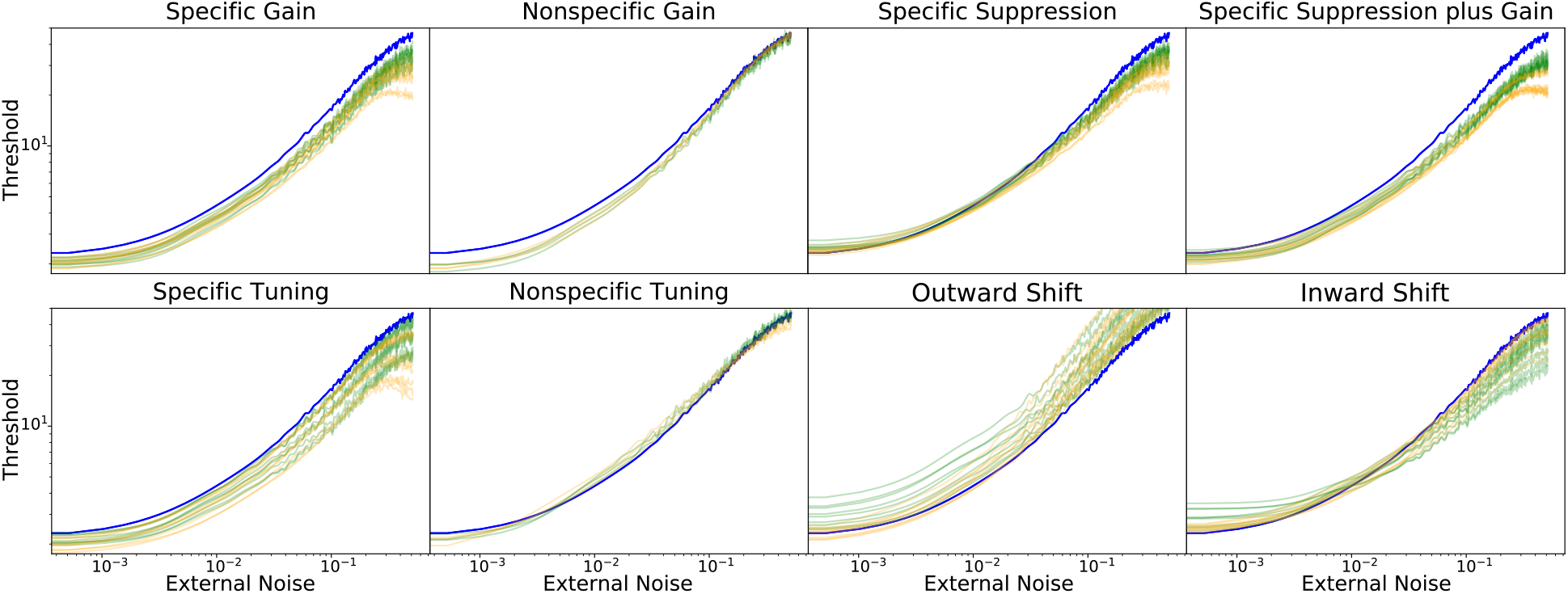
TvN curves are depicted for each mechanism of neural code change. Blue curves correspond to baseline models, orange curves to encoding changes implemented in a dense population (20 channels), and green curves to encoding changes implemented in a sparse population (10 channels).

From the point of view of the decoder, information about differences in stimulus values comes from differences in the responses of each channel. The slope of the tuning function at a particular value of the stimulus determines its sensitivity to changes in the stimulus (i.e., how different will be the response as a function of a small change in the stimulus). For a range of values, such slope can increase by increasing the height or decreasing the width of the tuning function.

Broadband noise involves adding random stimulation across all neural channels. Again, the slope of the tuning function determines to what extent small changes in random stimulation produce a large change in the channel’s response. In this case, however, a higher slope for channels other than those responding to a target stimulus means more influence of small variations in random noise, which decreases the precision of decoding for that target.

In “nonspecific” mechanisms of encoding change, the slope of tuning functions is affected equally across all channels. At low levels of external noise, this improves decoding precision, but at high levels of external noise, the effect of the noise on channels surrounding the target is stronger, worsening decoding precision. On the other hand, in “specific” mechanisms of encoding change, the slope of channels that respond to the target stimulus is affected more than the slope of surrounding channels that respond to noise, producing an overall improvement in performance across all noise levels.

The TvN curve produced after applying specific gain is consistently more precise (i.e., lower threshold) than that of the baseline across all levels of external noise. The increased responsiveness of channels around the target leads to steeper slopes in their tuning functions, This improves the channels’ sensitivity to stimulus changes around the target exclusively, without increasing sensitivity to stimulation due to external noise in off-target channels; thus, compared to baseline, the precision is improved across all levels of external noise. The TvN curve produced after applying nonspecific gain is initially more precise than that of the baseline during the flat and curved sections, but precision converges with the baseline by the end of the linearly-increasing section. During the flat section, much like specific gain, increased responsiveness leads to steeper slopes along channels surrounding the target, improving decoding precision. However, unlike specific gain, the responsiveness of the channels were indiscriminately increased; thus, during the linearly-increasing section, when the strength of the external noise becomes more comparable to that of the target stimulus, response to noise at the off-target channels is amplified and there isn’t an improved effect compared to baseline.

The TvN curve produced after applying specific suppression is higher than baseline (more imprecise) during the flat section, crosses baseline during the curved period, and it becomes lower than baseline (more precise) during the linearly-increasing section. The general effect of specific suppression is to decrease sensitivity to noise in the off-target channels. The level of suppression determines to what extent this also affects the channels close to the target. With high levels of suppression, the responsiveness of channels surrounding the target is reduced, which reduces their slope and precision of decoding at low levels of noise. The improvements in precision in the linearly-increasing section are due to the relatively stronger reduction of slopes in off-target channels, which makes them less sensitive to broadband noise. Figure 10 also shows an exception to the general pattern, in which specific suppression increases decoding precision in the flat part of the TvN curve. This happens when suppression is strong enough to reduce the influence of internal neural noise from channels surrounding the target, but weak enough to not disrupt the slope of those channels too strongly.

The TvN curve produced after applying specific suppression with nonspecific gain is generally more precise than that of the baseline across all levels of external noise. While the basic features are consistent with specific gain and specific tuning, the slope of the linearly-increasing section is more gradual along specific suppression’s TvN curve compared to baseline: both specific gain and tuning run in parallel to the baseline throughout the linearly-increasing section. Mechanistically, the improvement at high external noise levels is due to reduction of slopes in off-target channels, as in specific suppression. To this, non-specific gain adds an improvement at low external noise levels, due to an overall increase in slopes. Whether or not this improvement is enough to bring imprecision to a level lower than baseline depends on the balance between gain and suppression. As the nonspecific gain decreases, it becomes more similar to regular specific suppression; thus, the nonspecific gain is too low to represent a meaningful improvement from specific suppression. This explains some exceptions in Figure 10, in which imprecision is higher than baseline at the lowest levels of external noise.

The TvN curve produced after applying specific tuning is consistently more precise than that of the baseline across all levels of external noise. The TvN curves are seemingly identical to those produced after applying specific gain, and the underlying mechanism is again an increase in slopes for tuning functions surrounding the target. Due to the specificity of this mechanism, the change improves the channels’ sensitivity to stimulus changes around the target exclusively, without increasing sensitivity to stimulation due to external noise in off-target channels, producing a constant improvement in decoding along the linearly-increasing section of the TvN curve.

The TvN curve produced after applying nonspecific tuning is more precise than that of the baseline during the flat section, but decreases in precision during the curved section and the majority of the linearly-increasing section before finally matching the baseline. While the channels near the target have increased their sensitivity by aligning their steepest points to the target (explaining the flat section’s improvement), external noise is no longer uniquely beneficial at the target location as it was with specific tuning, and the indiscriminately improved sensitivity to stimuli (including external noise) is detrimental during the linearly-increasing section.

The TvN curve produced after applying an outward tuning shift is in most cases more imprecise than that of the baseline across all levels of external noise. In the general case, moving channels away from the target decreases the slopes of tuning functions at the target. This in turn decreases the channels’ sensitivity to stimuli near the target and reduces precision. The decreased channel sensitivity applies across all three sections of the curve: adding noise just makes it worse. The single exception among our simulations is one in which an improvement in target decoding is seen at low levels of external noise. We believe this to be an artifact of our simulation parameters. This effect is observed when the shift in tuning is so large that only a channel with the target as its preferred stimulus is left unshifted. The center of the tuning shift and the preferred stimulus of one channel must perfectly match for this to happen. Under such conditions, our assumption of a near-equal variance SDT model fails, because decoding precision is high exactly at the target (a maximum response value corresponds only to the target) but quickly decreases at stimulus values slightly off-target (due to symmetry in the tuning function, a given response value corresponds to two possible stimulus values).

The TvN curve produced after applying an inward tuning shift is generally more imprecise than that of the baseline during the flat and curved sections, but the precision improves during the linearly-increasing section (much like the pattern produced after applying specific suppression). During the linearly-increasing section, precision increases because the responses of the channels near the target are increased disproportionately by the high broadband noise compared to distal channels; furthermore, the coverage of the distal channels is decreased, reducing the effects that broadband noise can have there. During the flat section, imprecision of the target increases because the slopes of the shifted channels are flatter at the target.

Generally speaking, TvN curves are able to differentiate nonspecific gain, nonspecific tuning, and outward tuning shifts from all other models. Unfortunately, the TvN curves from inward tuning shifts are very similar to those of specific suppression, and populations affected by specific gain, specific tuning, and specific suppression plus nonspecific gain produce highly similar TvN curves as well.

### Threshold vs Stimulus (TvS) Curves

Differences in the number and properties of neurons encoding a particular dimensional value should produce differences in the precision with which that dimensional value can be decoded. In general, decoding from neurons which are more numerous, more finely tuned, and have a larger range of responses (i.e., difference between baseline and maximum firing rate) is more precise. For example, **(author?)** [6] showed that the precision with which orientation can be decoded from a neural population strongly depends on the number of cells encoding such variable. Due to the cortical magnification factor in V1, the number of cells encoding orientation drops with eccentricity, and this drop provides an excellent fit to estimates of orientation thresholds as a function of eccentricity.

The different mechanisms of change in population codes shown in Figure 3 should therefore produce concomitant changes in thresholds measured at different values of the stimulus dimension. An experiment measuring thresholds at many different “pedestal” values of the dimension should produce a Threshold versus Stimulus (TvS) curve that would provide important information about underlying changes in population codes. A TvS curve is also much easier to interpret than many other possible psychophysical functions. Sections of the curve with higher values represent more imprecise decoding estimates, which result from neurons that are relatively fewer in number, more broadly tuned, or with a smaller response range. Sections of the curve with lower values represent more precise decoding estimates, which result from neurons that are relatively more in number, more finely tuned, or with a larger response range.

As shown in Figure 11, the sections of the TvS function are loosely categorized based on their distance from the target. Given that our simulations focus on circular dimensions, such as orientation, stimuli farthest from the target are shown on the left and right side: the distal section. The target itself and nearby stimuli fall under the target section. The remaining cases fall under the intermediate section. The TvS functions were all acquired in the absence of external noise, which means the target section should correspond to the initial flat region of the TvN curves.

**Figure 11:**
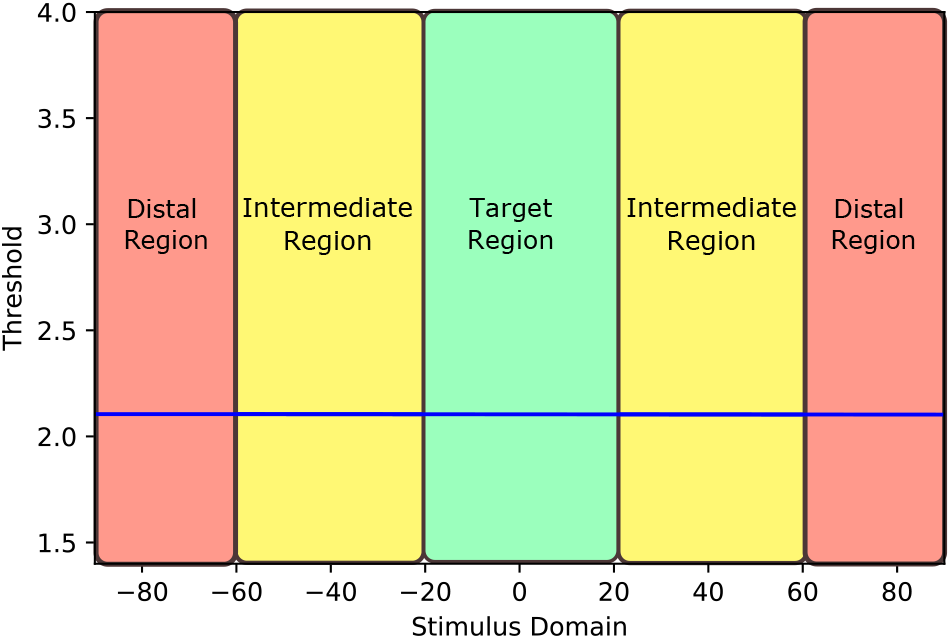
Division of the stimulus dimension into regions as a function of their distance from the target stimulus. The division is rather arbitrary, and we propose it here mostly as a way to interpret the simulated TvS curves shown in Figure 12.

The TvS function produced after applying specific gain is generally more precise than that of the baseline, but gradually approaches the baseline precision as the effect dissipates toward the distal section. The sensitivity of the channels are increased most in the target region (due to increased slopes of tuning functions in that region) and least in the distal region, explaining the effect.

The TvS function produced by applying nonspecific gain is always more precise than that of the baseline. The channels’ slopes are increased indiscriminately, allowing the effect observed in specific gain to be applied across the stimulus domain. The level of gain directly determines the degree of the increase in decoding precision. Without the interference of external noise, the increase in responsiveness is useful for all values of the stimulus domain.

The TvS function produced by applying specific suppression is always more imprecise than that of the baseline. The imprecision increases toward the distal sections. Specific suppression produces flatter slopes across tuning functions of all channels, reducing decoding precision, but the effect is stronger the farther away a channel is from the target.

The TvS function produced by applying specific suppression plus gain is more precise than the baseline at the target and intermediate sections, but more imprecise in the distal section. The shape of the TvS function matches specific suppression exactly, but its vertical position is determined by the level of nonspecific gain: adding enough nonspecific gain can lead to a TvS curve exactly like that of specific gain, and adding too little leads to a TvS curve exactly like that of specific suppression. However, if enough gain was modeled to match or surpass baseline along all sections, the mechanism would not truly qualify as involving suppression. The TvS function produced after applying specific tuning is more precise than baseline in the target section. Within the target section especially, the slopes of tuning functions are increased, which accounts for the corresponding improvements in precision. In 50% of the simulations, specific tuning caused the TvS function to increase imprecision above the baseline during the intermediate and/or distal sections. Finding such “shoulders” in the TvS function would differentiate specific tuning from specific gain, but not finding them is insufficient to differentiate this mechanism from specific gain (this comparison is particularly important, as TvN curves for these two models are also indistinguishable). For the rest of the simulations, five of them (27.8%) involved a change in tuning so small that no change was visible in the TvS curve except for a very small increase in performance for all values of the stimulus. The last three cases involved curves similar to those observed for specific gain. Thus, there are cases in which the TvS function for specific tuning mimics that observed for specific gain.

The TvS function produced after applying nonspecific tuning is more precise than baseline across all sections. As another flat line, the function is seemingly identical to the nonspecific gain function. The uniform precision increments are due to the indiscriminate increase in slopes across tuning functions. As shown in the corresponding TvN curve, the improvement only applies without the interference of external noise.

The TvS function produced after applying an outward tuning shift is generally less precise than baseline at the target section, and the precision generally improves at the distal section. Shifting of tuning functions away from the target tends to concentrate slopes in the farthest area, where more precise decoding is made possible. The intermediate channels, which move the most, also cannot assist with decoding stimuli in the target region. The redistribution of slopes seems beneficial to decoding only starting at intermediate distances from the target (i.e., around +/−50), but is detrimental to decoding at values closer to the target. The TvS function produced after applying an inward tuning shift is less precise at the target region, more precise in the intermediate region, and again less precise in regions farthest from the target. Shifting of tuning functions towards the target tends to concentrate slopes in the intermediate area, with the consequence of a lower concentration of slopes in the region closest to and farthest from the target.

Generally speaking, TvS functions are able to differentiate specific suppression, specific suppression plus gain (when parameters do not make either of the two mechanisms dominate), outward shift, and inward shift from all other models. Unfortunately, the TvS functions for specific gain and tuning were too similar to each other to distinguish them, and the functions for nonspecific gain and nonspecific tuning were practically identical. However, TvS functions allow researchers to still narrow down the type of change in encoding to either a specific or nonspecific mechanism. If a nonspecific mechanism is found, then gain and tuning can be further separated by using TvN functions, as shown in the previous section.

## Discussion

We performed a large number of simulations to determine what types of neural encoding changes could be differentiated through psychophysical threshold experiments. A summary with the most common patterns of results observed in our simulations is shown in Table 1. The results suggest that, by gathering thresholds along the stimulus domain (i.e., TvS curves), it is possible to distinguish four of the eight types from all other mechanisms: specific suppression, specific suppression plus gain (as long as both mechanisms have a balanced contribution), outward tuning shift, and inward tuning shift. From the remaining four mechanisms, the pair of nonspecific mechanisms (gain and tuning) can be distinguished from the pair of specific mechanisms (gain and tuning) by how much thresholds for stimuli outside the target area are affected. If the candidate mechanisms are nonspecific, then an additional experiment that manipulates external noise (i.e. TvN curves) could help finalize the selection. If the candidate mechanisms are specific, then an increase in thresholds for intermediate and/or distal regions of the TvS curve is indicative of a specific tuning mechanism. In a minority of cases (3/18 of the simulations), a specific tuning mechanism produced a TvS curve mimicking that of a specific gain mechanism, and thus both cannot be separated unless other sources of information are taken into account (e.g., inverted encoding modeling).

**Table 1:**
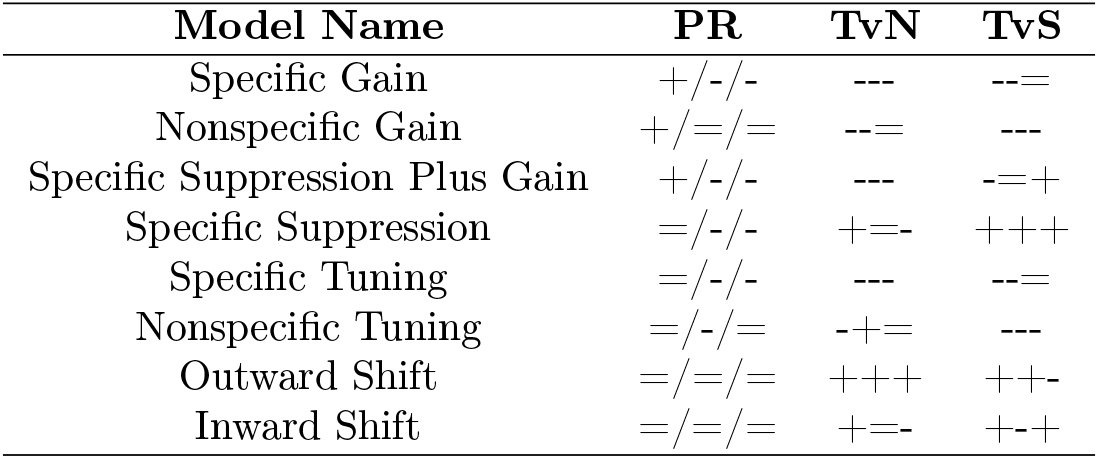
Most common patterns of results observed in our simulations, with symbols representing changes from baseline. Values higher, equal, and lower than baseline are represented by +, =, and -, respectively. Symbols in the Population Response (PR) column represent difference with baseline in the three parameters of a curve fitted to the population response to the target: curve height, curve width, and peak curvature, in that order. Symbols in the Threshold versus Noise (TvN) column represent difference with baseline in the flat, curved, and linearly increasing sections of the curve, in that order. Symbols in the Threshold versus Stimulus (TvS) column represent difference with baseline in the area around the target, in the intermediate section, and the distal section, in that order.

While TvN and TvS alone could not dissociate every mechanism (i.e., specific gain and specific tuning produced qualitatively identical patterns), the addition of other data may allow to infer the true mechanism of encoding change. For example, it is possible to estimate population responses from neuroimaging data using inverted encoding modeling. In addition, additional psychophysical studies may be able to differentiate between the two mechanisms, as they should differ in the overall level of activity produced by presenting the target. Assuming that the overall level of activity elicited by a stimulus determines its relative salience, producing bottom-up attention, then a target under specific gain should capture attention more easily in visual search tasks than a target under specific tuning [43]. More generally, at the heart of our approach is the idea that no single study allows one to infer the correct mechanism of encoding change underlying some behavior of interest. Rather, a combination of multiple studies, all linked together through the same model, provides a much stronger approach to the problem.

A number of assumptions were made in this simulation work, which are listed in what we perceive is their order of importance in Table 2. First, because there is inherent ambiguity in linking the neural and psychophysical levels, such that any changes in thresholds could be attributable to either encoding or decoding changes [23], we started by assuming an optimal decoder. This is a rather strong assumption, but is also standard in the prior literature [e.g., 1, 24, 3, 4, 7, 6], where it has proven to be useful. In addition, biologically plausible mechanisms for optimal decoding in the brain have been proposed in the literature [24]. Second, we also assumed independent Poisson neural noise, which facilitates maximum likelihood estimation and is also a common assumption in the literature [e.g., 1, 24, 3, 4, 7, 6]. The strongest part of this assumption is that noise is independent, although we have no reason to expect that correlated noise would change our general conclusions. Third, we assumed that decoding noise is similar in neighboring areas of the stimulus dimension. Once again, this is a common assumption in the literature linking neural encoding with psychophysics [1, 3], made mostly for convenience as it substantially reduces the computational cost of simulations. We believe that this assumption also seems valid, as there is little reason to expect large differences in decoding precision in a small area of the stimulus domain. Fourth, we assumed a bell-shaped tuning function, which as far as we know is the only type of function used in research applying inverted encoding modeling [e.g., 9, 38, 31, 15, 13, 11, 12, 10]. For other stimulus dimensions, such as those characterized by monotonically increasing or decreasing tuning functions (e.g., face dimensions; [44, 5, 45, 43]), new simulations would be required. Finally, we assumed a homogeneous encoding population for the baseline condition. This is a common assumption in inverted encoding modeling, and we don’t think that changes in this assumption would change any of our conclusions. This is because, regardless of what baseline is assumed, changes in encoding relative to that baseline should result in similar changes in decoding precision relative to the baseline.

**Table 2:** List of assumptions made in the present simulation work, ordered from strongest to weakest.

| Assumptions |
| --- |
| Optimal MLE Decoding |
| Independent Poisson Neural Noise |
| Equivalent Noise Variance in Neighboring Areas of Stimulus Dimension |
| Bell-Shaped Tuning Function |
| Homogeneous Encoding Baseline* |

As we have indicated, some of the assumptions in Table 2 are rather strong. However, we do not believe that they are stronger than the assumptions implicit to inverted encoding modeling, discussed in the introduction section. Again, we believe that the weaknesses of each approach can be overcome by a research approach that combines different sources of data (which require different sets of assumptions) within a single modeling framework.

### Re-interpreting results in the literature

As indicated in the introduction, we have focused here on those mechanisms of encoding change that have been proposed to improve task performance in the previous literature, making them candidates to explain the effects of learning and attention on perceptual processing. Although no prior experiment has attempted to distinguish between multiple mechanisms using both TvS and TvN functions, several experiments have gathered thresholds either at the target stimulus (i.e., the stimulus involved in learning or attention), across different values of the stimulus dimension (i.e., similar to a TvS function), or across different levels of external noise (i.e., a TvN function). The results of some of these studies can be re-interpreted in the light of our current simulations.

For example, several studies have shown that aversive Pavlovian conditioning involving a particular stimulus (the conditioned stimulus, or CS+, which is paired with an aversive stimulus, such as electric shock) produce increments in thresholds for that stimulus [46, 47, 29]. Figure 12 shows that only a few mechanisms can produce this increase in thresholds at the target: specific suppression, outward shift, or inward shift. However, other evidence suggest that, among these candidates, the most likely mechanism is one of inward shift. First, neurophysiological studies in auditory aversive conditioning with rodents have shown evidence that individual neurons shift their preferred stimulus towards the CS+ after training (for reviews, see [39, 40]). Second, there are multiple reports that an aversive CS+ captures attention in search tasks [48, 49, 50, 51]. A reasonable assumption is that bottom-up attentional capture depends on the overall level of neural activity that the CS+ produces in comparison with concurrently-presented stimuli that compete for attention [43]. If we think of that overall level of neural activity as the result of both the number of neurons selective for the CS+ as well as their firing rates, we see that both suppression and outward shift are mechanisms likely to reduce the overall level of neural activity produced by the CS+, whereas an inward shift is likely to produce the opposite effect. In sum, both neurophysiological and psychophysical data suggest that the most likely change in stimulus encoding produced by aversive conditioning is an inward shift towards the CS+. This hypothesis could be easily tested by estimating TvS functions from participants before and after conditioning. Based on our results, we would predict that the post-learning TvS function would look like the inward shift function in Figure 12.

**Figure 12:**
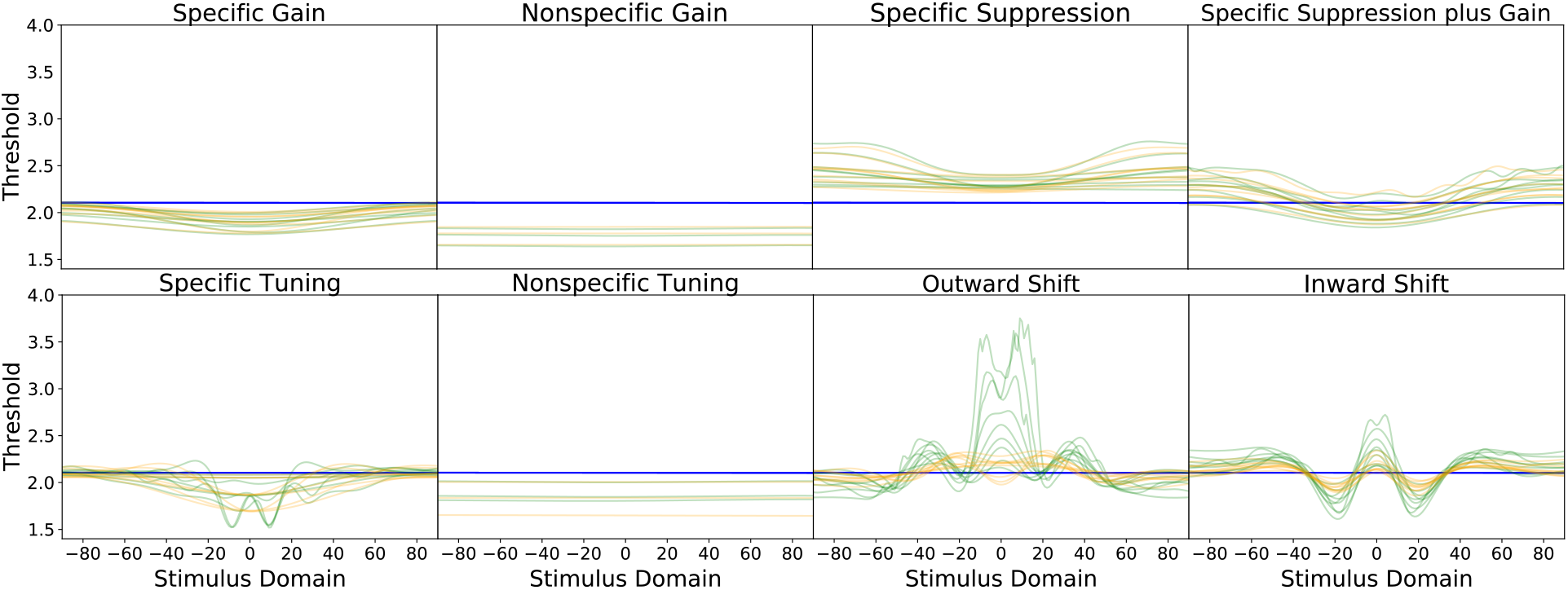
TvS curves are depicted for each mechanism of neural code change. Blue curves correspond to baseline models, orange curves to encoding changes implemented in a dense population (20 channels), and green curves to encoding changes implemented in a sparse population (10 channels).

Another form of learning known to produce changes in dimension discriminability is category learning, although such changes have usually been measured using a measure of sensitivity (i.e., *d*′) rather than sensitivity thresholds. In particular, several studies have shown that categorization training produces increased discriminability along the category-relevant stimulus dimension [52, 53, 54, 55, 56, 57, 58], an effect that can be sometimes stronger for stimuli that cross the category boundary [55, 56]. On the other hand, a recent study using inverted encoding modeling [38] has found evidence supporting the operation of an outward tuning shift mechanism during categorization of oriented gratings. These two results are difficult to interpret together. In particular, Figure 12 shows that the outward shift mechanism found by Ester et al. in most cases would lead to a reduction in stimulus discriminability (i.e., increased thresholds, or decoding imprecision) around the bound (which in this case corresponds to the target at zero). Improvements in discriminability should be reliably observed only for stimuli that are relatively far from the bound. In line with this idea, Van Gulick et al. [58] report some evidence of a stronger effect on discriminability away from the category bound than at the category bound. On the other hand, Goldstone [55] reports the opposite pattern of results, and others [52, 53] report equivalent improvements in discriminability across the dimension. What all these studies have in common is that they report an increment in discriminability for stimuli around the category bound, which is difficult to explain by an outward shift mechanism.

It is possible that a precisely-tuned outward shift can produce this result, if the shifts place the slopes of several tuning functions around zero. In any case, an outward shift mechanism predicts a loss in discriminability in some areas of the dimension, which has not yet been observed. Because a tuning shift necessarily increases discriminability in some areas at the cost of a reduction in discriminability in other areas, we predict that if an outward shift is the mechanism underlying perceptual effects of categorization, then a fine-grained TvS curve should reveal areas in which the category-relevant dimension shows a reduction in stimulus discriminability.

Another possibility is that the precise mechanism of encoding change produced by categorization learning is dependent on properties of the neural population encoding a particular stimulus dimension. More precisely, Ester et al. [38] used orientation of lines and gratings as their stimuli, and orientation is known to be encoded in early visual cortex through tuning functions similar to those used in the present study. On the other hand, the majority of the psychophysical research has used either highly complex shape and object stimuli [52, 53, 54, 57, 58] or simple dimensions other than orientation [55, 56]. The tuning functions used by the brain to encode such dimensions might be different than what is represented by the standard population model used here. For example, face features are thought to be encoded through monotonic tuning functions (e.g., sigmoidal; see [5, 45, 43]). Using computational modeling and visual adaptation, it has been found that the effects of categorization on perception of face identities along the category-relevant dimension [59, 60, 61] can be best explained using a specific gain mechanism [45]. It is currently unknown exactly how the complex shape and object stimuli used in some studies are encoded, but encoding that is different from that of orientation might be at the heart of the results obtained with such dimensions.

Other studies have reported TvN curves to characterize the effects of learning in terms of psychophysical observer models [62]. The results of such studies can be re-interpreted in terms of the encoding/decoding observer model studied here (Figure 1). For example, perceptual learning results in TvN curves that drop from baseline at all levels of external noise [63, 64, 65]. This result is consistent with multiple mechanisms of encoding change (see Figure 10): specific gain, specific tuning, and specific suppression plus gain. Interestingly, all these mechanisms are in line with the well-known stimulus specificity of perceptual learning. To distinguish among these different potential mechanisms, more information can be obtained through a TvS curve, but additional steps might be required to differentiate between specific gain and tuning (see section *Recommendations for researchers* below).

Finally, There are multiple studies that have estimated TvN curves under different attentional demands. The study that is closest to our work was performed by Ling et al. [**?**], who specifically sought out to dissociate between nonspecific gain and specific suppression with nonspecific gain as mechanisms of attention. The authors chose those two mechanisms because they would specifically reduce thresholds in the early (internal noise suppression) and late (external noise suppression) parts of the TvN curve, respectively. Our simulations have confirmed that, among all the mechanisms of encoding change studied, these two are the only ones that seem to uniquely affect internal or external noise. Thus, our results support the methodology used by Ling et al. [**?**], although it must be noted that specific suppression plus gain must be finely tuned in order to produce no effect in the early part of the TvN curve. In most cases, the TvN curve drops below baseline at low levels of external noise (see Figure 10).

The results reported by Ling et al. suggest that the mechanism of encoding change underlying spatial attention is nonspecific gain, which is in line with other reports [66]. On the other hand, feature-based attention produced a drop in the TvN function at all levels of external noise, which as mentioned earlier is consistent with multiple mechanisms. None of those mechanisms were considered as plausible hypotheses by Ling et al., who instead preferred to interpret their results as suggesting a combination of specific suppression with nonspecific gain (“tuning”), and an independent mechanism of nonspecific gain (“gain”). We believe that this combination of mechanisms is a less parsimonious explanation for their results than any of the models producing an overall drop in the TvN curve shown in Figure 10.

An important note of caution is necessary when reinterpreting some TvN results from the attention literature. In many cases, the TvN curves presented do not represent the presence and absence (i.e., baseline) of an attentional effect. Rather, they represent a comparison between two different attention effects, such as valid versus invalid cues [67] or pre-cueing versus simultaneous cueing [68]. Such results are more difficult to interpret than the simpler case in which a baseline (i.e., absence of any task feature or experimental manipulation) is compared against a simple manipulation (i.e., the presence of a task feature or experimental manipulation), and the simulations presented here can only provide information about this latter, simpler case.

### Recommendations for researchers

When measuring participants’ psychophysical thresholds, we recommend using sensitivity thresholds (a stimulus value related to a specific value of *d*′) rather than the more commonly-used accuracy thresholds (a stimulus value related to a specific proportion of correct responses). While accuracy thresholds would be contaminated with bias, sensitivity thresholds take response bias into account. There are currently methods available to estimate sensitivity thresholds using both yes/no and 2AFC tasks [69]. An important theoretical reason to prefer sensitivity thresholds is that the theory links them directly to decoding precision in the encoder/decoder observer model.

Also, we found that obtaining estimates at the target stimulus, with or without the inclusion of external noise (i.e., TvN curve), is insufficient to distinguish between all of the mechanisms of encoding change. Thus, we recommend gathering estimates for stimuli surrounding the target as well (i.e., TvS curve), to elucidate the neural mechanisms at work.

No singular experiment could distinguish every possible change in neural codes. We recommend acquiring thresholds as a function of multiple factors throughout multiple experiments, but if manipulating only one factor is possible, then measuring thresholds along the stimulus domain seems to be very effective (showing unique patterns in all cases, except to distinguish some cases of gain versus tuning). On the other hand, if a researcher is interested in telling apart a specific mechanism of encoding change from all others, then it could be wiser to obtain a TvN curve instead. For example, either nonspecific gain or nonspecific tuning can be separated from all other mechanisms using a TvN curve, but not a TvS curve. When faced with uncertainty between two candidate models, a good approach might be to fit the models to the observed data and perform formal model selection [70, 71, 72], although this would be a computationally intensive procedure employing the Monte Carlo technique used here to obtain thresholds from the model.

It is also important to note that these models were differentiated under specific assumptions about encoding and decoding; therefore, we recommend that researchers carefully consider what assumptions they will be able to justify (i.e., adaptation researchers cannot justify the use of an “aware” optimal decoder; see [7]). However, we must note that the assumptions necessary to make inferences about mechanisms of encoding change from psychophysics do not seem stronger than equivalent assumptions necessary for the application of inverted encoding modeling of neuroimaging data (normal neural noise, additive independent measurement noise, linear measurement model).

Here, we have focused mostly on basic mechanisms of encoding change, meaning mechanisms that alter a single aspect of the population of tuning curves. Caution should be maintained when combining basic mechanisms into more complex ones. In our simulations, specific suppression plus gain was consistently an interpolation between specific suppression and specific gain. If it is necessary to test models implementing combinations of these mechanisms of encoding change, the parameters need to be balanced to avoid ambiguity. In our example, specific suppression plus a very small gain looks very similar to ordinary specific suppression, whereas large gain with small specific suppression looks very similar to ordinary nonspecific gain.

In the future, a powerful methodology would involve using both psychophysics and neuroimaging data together to infer changes in encoding, perhaps using hierarchical Bayesian modeling, in which multiple types/sources of data for each participant can be used simultaneously to make inferences about a single set of model parameters [73].

### A warning regarding “behavioral tuning curves” obtained through pattern masking

Pattern masking experiments involve the presentation of a target stimulus combined with a pattern stimulus that “masks” or interferes with the perception of the target [74]. It is possible to measure sensitivity to a constant target as a function of changes in some property of the mask pattern. Sensitivity is measured via thresholds, which as before estimate the imprecision of target decoding. Figure 9 shows a sketch of the typically-found function, which here we will call a Threshold versus Pattern Masking (TvP) curve. As shown in the figure, the typical pattern is one in which masking becomes greater (i.e., thresholds or imprecision becomes higher) as the mask becomes closer to the target in orientation (or any other feature).

The TvP curve is thought to reveal the shape of a “psychophysical filter” used to detect the target [25, 74]. The width of the non-flat part of the curve (i.e., blue area in Figure 13) represents the area where the mask interacts with such filter, and it can be widened or narrowed [27]. The height of the curve may also change, both across all values of the pattern mask [25, 37, 27] as well as in specific values [25], representing corresponding increments or decrements in the amount of interaction between a mask and the target.

**Figure 13:**
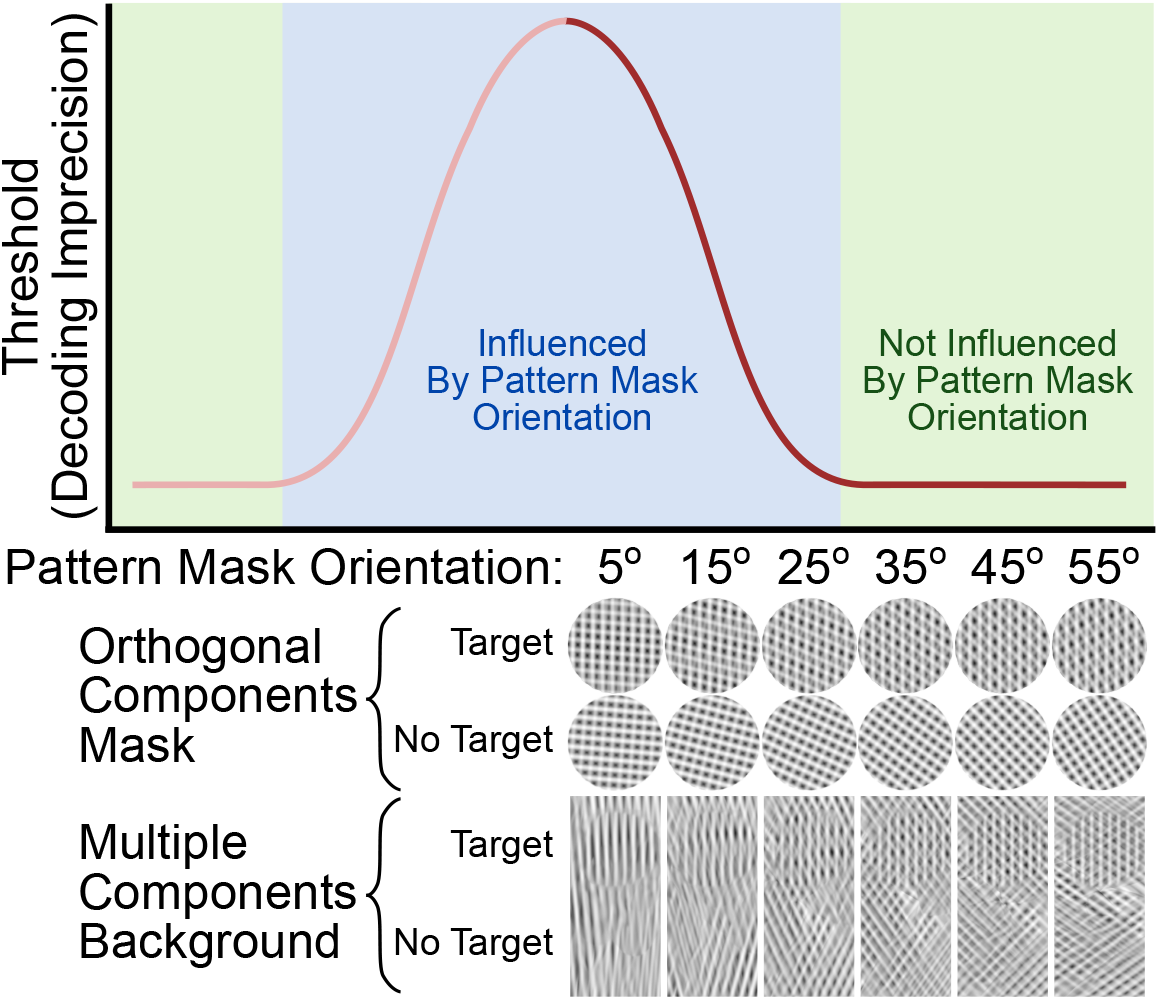
Explanation of a Threshold versus Pattern Mask (TvP) curve.

TvP curves resemble tuning functions of neurons in visual cortex, and their bandwidth may also reflect properties of such tuning functions [26]. For this reason, many researchers interpret them as if they represent a form of “behavioral tuning curve” that can be directly linked to neural mechanisms of change in population codes [e.g., 25, 26]. However, it is important to underscore that TvP curves do not provide estimates of either neural tuning functions or of population responses such as those shown in Figure 1. In fact, we will argue that TvP curves cannot be studied using the theoretical framework used here.

Indeed, the likely mechanisms behind pattern masking experiments go beyond the standard population encoding model that is the focus of the present and prior research (see Figure 1). TvP curves can be obtained using at least two procedures. The first is to use a mask composed of two orthogonal components, and ask participants to discriminate between the mask presenting a target versus the mask presenting no target (see Orthogonal Components Mask in Figure 13; [25]). In V1, the effect of superimposing a mask grating on top of a stimulus with the neuron’s preferred orientation is a reduction in responding known as *cross-orientation suppression* [75, 76], and its mechanism is thought to involve interactions between several neurons with overlapping receptive fields [75, 77]. In the simple standard population encoding model, the responses of neural channels are independent from one another. This facilitates simulation in simple conditions, but means that the cross-orientation suppression present in pattern masking experiments cannot be simulated without a substantially more complex model [e.g., 2].

A second procedure used to obtain a TvP curve is to use a large background composed of multiple stimulus components, embed the target somewhere in this background, and ask participants to indicate where the target has been presented (see Multiple Components Background in Figure 13; [37, 27]). Because the target has some level of transparency, cross-orientation suppression is also involved in this type of experiment. In addition, we can expect the presence of orientations surrounding the target to produce additional effects, which can be either suppressive or facilitatory in nature [78]. That is, this second type of experiment, involving cross-orientation suppression as well as surround suppression/facilitation, is absolutely beyond the scope of the simple standard population encoding model.

The previous considerations suggest that the TvP is not an estimate of either classical neural channel tuning functions or population responses, and simple interpretations of the TvP curve in those terms [e.g., 25, 26] are not appropriate. In line with this idea, we attempted simulating TvP curves in many different ways using the framework presented here, but our simulations never captured even the basic shape of the curve. The reason is simple: as a mask that only includes stimulation within a certain range of stimuli is moved closer to the target stimulus, the likelihood of the target and the precision of decoding *increases* (i.e., the opposite of the pattern of results shown in Figure 13). That is, the mask provides a boost in activity at the target, similar to a mechanism of specific gain. The model cannot capture the interference between mask and target, because such interference involves mechanisms beyond the classical tuning functions represented in the standard population model.

### What about other encoding and decoding models?

For orientation stimuli, we chose to model channels as symmetrical Gaussians with Poisson noise based on previous literature, but there are other ways to characterize tuning functions that may be more appropriate for other types of stimuli. For example, face dimensions are often modeled using asymmetric sigmoidal tuning functions [44, 5, 45, 43]. Also, another common assumption for neural channel noise is that it follows a Gaussian distribution [e.g., 6, 7], a common assumption implicit in inverted encoding modeling analyses of neuroimaging data [see 8, 14].

In addition, we chose an optimal decoder that would be considered “aware” [7]. That is, we assumed that the decoder had complete knowledge of the statistics of each encoding model, before and after application of a given mechanism of encoding change. It is known that this assumption is unlikely to hold for short-term neural code changes, such as those underlying adaptation [7]. However, we think that an aware decoder provides a better assumption for situations involving encoding changes that are predictable due to extensive prior experience, such as those produced by learning and other cognitive mechanisms.

In addition, although optimal decoding through MLE has been widely used, other decoding schemes are possible. For example, it is possible to use simpler linear decoders, or if the population’s channels are dense enough, a max response decoder. However, MLE is biologically plausible [24] and provides a more meaningful benchmark due to it being optimal for the task.

### Conclusion

Psychophysical thresholds can help researchers infer the mechanisms of encoding change due to cognitive states of interest. Together, the TvS and TvN functions can differentiate six of the eight mechanisms under study here (in some cases, seven). Assessing the decoding precision through thresholds is a useful tool to supplement inverted encoding models obtained from neuroimaging to estimate the population response. Because any quantifiable stimulus dimension can be used to produce a TvS or TvN function, this approach can be applied to a variety of high- and low-level stimuli to answer a plethora of questions in vision neuroscience.

## Methods

### The encoding-decoding observer model

In the computational neuroscience literature, population encoding models (see Figure 1) are used to formally describe how populations of neurons represent a stimulus through their patterns of activity [17, 16, 79]. In the case of stimulus dimensions, an encoding model represents how changes in a dimension of interest are related to changes in neural responses. The response of each neural channel (i.e., neuron or population of neurons with similar properties) can be described by a tuning function. A commonly chosen tuning function *f*_*c*_(*s*) to represent the mean response of each channel (*c*) to any presented stimulus value (*s*) is a Gaussian curve, where 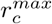 represents the maximum height of the channel, *ω*_*c*_ represents the the standard deviation of the curve (i.e., width), and *s*_*c*_ represents the channel’s preferred stimulus:

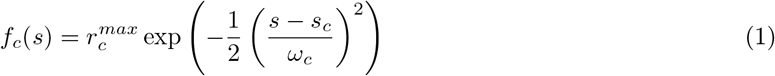

However, channels’ responses can be influenced by neural noise, which here we describe using a Poisson distribution, independently for each channel [17, 18].

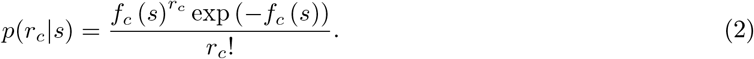

When a stimulus with a given value in the dimension *s* is presented to the model, the output of the population encoding model is a vector of neural responses **r**, which implicitly encodes information about that stimulus value. As in previous research [e.g., 1, 24, 3, 4, 7, 6] we assume that such information is recovered through optimal decoding via maximum likelihood estimation (MLE).

Generally speaking, the ML estimate is obtained by finding the value of *s* that maximizes the probability of the population response (**r**) given a fixed encoding model:

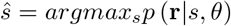
where *θ* represents a vector with all the fixed model parameters. When neural noise is independent, 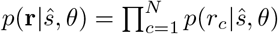, and the sum of the logarithms can be maximized instead:

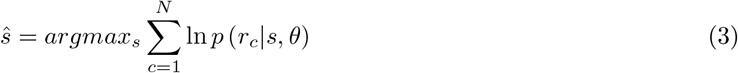

For a model with independent Poisson noise and Gaussian tuning functions, when homogeneous populations are involved, there are closed-form expressions [80] as well as simple algorithms [17] to solve Equation 3. However, several of our simulations involve non-homogeneous populations, and thus we obtained optimally decoded stimuli by finding the value of *s* that m aximizes the f ollowing equation:

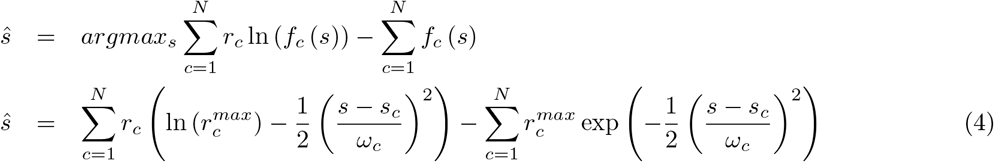

Numerical maximization of equation 4 requires providing a starting value for the optimization algorithm. We started the optimization at *s*, to increase speed and precision, as we know that *ŝ* must be in the vicinity of the true stimulus value *s*.

A property of MLE is that as the number of neural channels increases, the distribution of the decoded stimulus estimate becomes well-approximated by a Gaussian distribution with a mean of *s* and variance equal to:

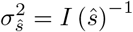
where *I*(*ŝ*) is the Fisher information at the ML estimate. For complex, non-homogeneous encoding populations, 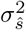 can be estimated through Monte Carlo simulations [1, 3]. At each repetition *m* = 1, 2, …*M*, the model is presented with a given stimulus and the noisy population response is used to decode an estimate of the stimulus value 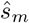. Using all *M* repetitions, one can obtain an estimate of the variance of the distribution of *ŝ* through the following equation:

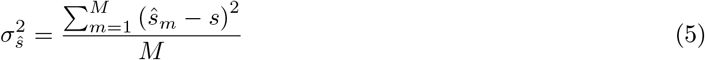

### Link to sensitivity thresholds

The encoding/decoding observer model presented in the previous section can be linked to signal detection theory (SDT) if one assumes that the decision variable in SDT is determined by decoding a population response. The decoded stimulus value *ŝ* corresponds to SDT’s decision variable, which is compared to a decision criterion (*c*) to produce a decision. Because the MLE estimate *ŝ* follows a Gaussian distribution with mean *s* and variance 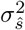, one can write the following definition of sensitivity:

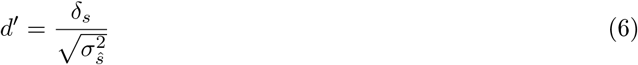
where *δ*_*s*_ refers to a small change in the dimension required to detect a change in the target stimulus *s* with a sensitivity equal to *d*′. Note that this difference is small enough that we assume an SDT model with a common noise variance for both stimuli, 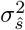. Re-arranging Equation 6 provides an equation to obtain a sensitivity threshold *δ*_*s*_ associated with a given value of *d*′:

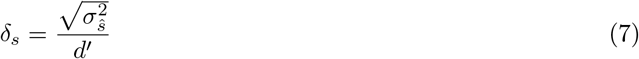

In our simulations, we use *d*′ = 1, a case in which thresholds are equivalent to the standard deviation of MLE estimates around *s* (see Figure 1b).

### Model Variations Under Study

All our simulations follow a similar pattern: starting with a homogeneous baseline set of encoding channels, we generated several variants of the encoding model, each representing a different mechanism of encoding change, and simulated psychophysical functions involving sensitivity thresholds obtained through Equation 7). In this section, we describe how we obtained all the tested population encoding models, and later discuss the simulation procedures in more detail.

To focus on changes in the psychophysical functions that were reliably produced by each mechanism of encoding change, several versions of each model were simulated, and they differed in the magnitude of the effect on encoding and/or the concentration of the effect around the target stimulus. Three effect magnitudes were simulated for each model, and the parameters used to obtain them are summarized in Table 3 and described in more detail below for each model. In addition, for all mechanisms of encoding change classified as “specific,” meaning that their effect was strongest at the target stimulus and gradually decreased with distance from the target, we included three additional variants that differed on the concentration of the effect around the target stimulus. In those cases, the “magnitude” variants were factorially combined with the “concentration” variants, to obtain 9 possible versions of each model.

**Table 3:**
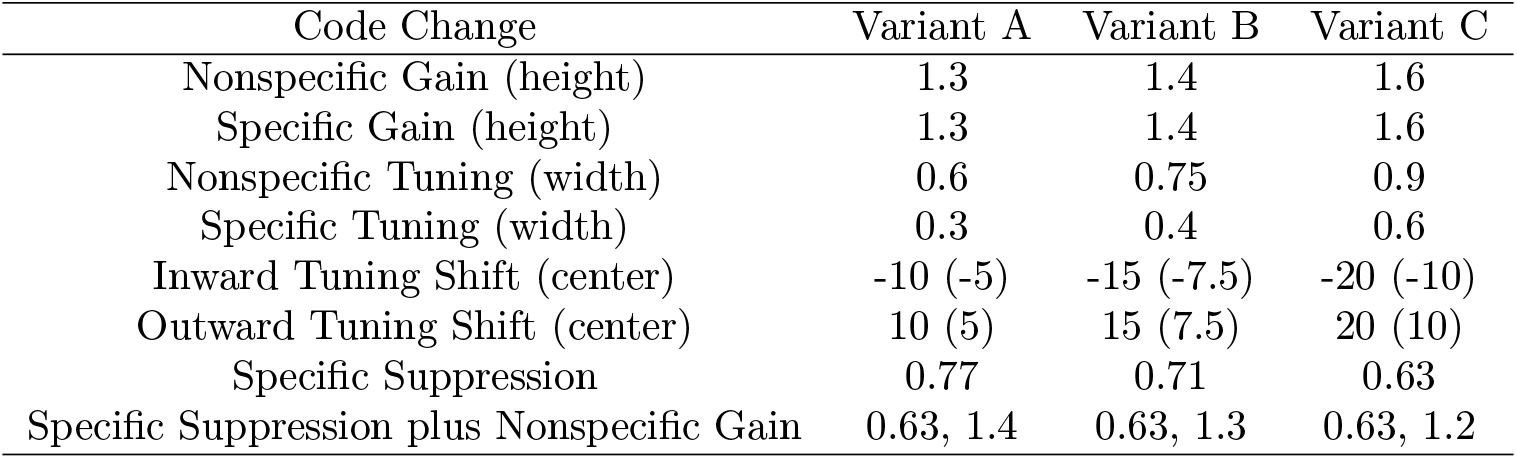
Parameters used to obtain different effect magnitude for each mechanism of encoding change under study. In addition, every “Specific” variation, including the tuning shifts, had 3 additional variants where the effect falloff was increased (1.0, .667, .333—.333 caused the effect to falloff most quickly, 1.0 caused the effect to falloff at the ‘edge’ of the circular domain—the farthest points from the target). For tuning shifts, the dense populations had reduced parameters (in parentheses) to prevent channels from crossing/overshooting the target. Specific suppression was implemented as a combination of Specific Gain and Nonspecific Gain.

As shown in Table 3, we studied 8 different mechanisms of encoding change: specific/nonspecific multiplicative scaling, specific/nonspecific bandwidth narrowing, inward/outward tuning shifts, as well as specific suppression with and without nonspecific multiplicative scaling (the latter being the only combination of two basic mechanisms, included because it was suggested in the prior literature as a mechanism involved in selective visual attention [3]). For an explanation of why these specific mechanisms were selected, see the Introduction section.

Mechanisms of encoding change categorized as “specific” (including tuning shifts) had a target stimulus for which the effect was strongest, and from which the effect gradually spread to other dimensional values. In our simulations, this target stimulus was always at the center of the dimension, which was labeled a zero degrees. The strength of each specific effect decayed linearly with the distance from the target, starting at the target and ending at 1, 2/3, or 1/3 of the half-range of the stimulus dimension. We call this value the *scaling width*, and it represents the inverse of the concentration of the effect around the target. The closer the scaling width is to 1, the more spread out the effect is towards the end points of the dimension. On the other hand, nonspecific changes apply evenly across the domain.

#### The baseline homogeneous Population

The baseline for all of the simulations was a homogeneous population of channels with *ω*_*c*_ = 12 and *r*_*max*_ = 12. We assumed a circular stimulus domain varying in degrees, as with grating orientation, so the channels placement ranged from [−90,90) (note, however, that multiple other stimulus dimensions are described by such circular domains). There were two versions of the baseline homogeneous population, and all the encoding changes specified below were applied to each model. The sparse baseline had 10 channels, placing the channels’ centers *s*_*c*_ at −90, −72, −54, −36, −18, 0, 18, 36, 54, and 72. The dense baseline had 20 channels, placing the channels’ centers *s*_*c*_ at −90, −81, −72, −63, −54, −45, −36, −27, −18, −9, 0, 9, 18, 27, 36, 45, 54, 63, 72, and 81.

#### Nonspecific gain

If the height of the tuning functions is uniformly altered by a scaling factor across the entire stimulus domain, then the change is referred to as nonspecific gain (see Figure 3). The changes in height directly correspond to changes in the average maximum responsiveness for the channels (i.e., multiplication by *r*_*max*_ at all channels). The scaling factors we tested were all above 1 because increases in responsiveness are expected to improve thresholds; thus, we tested: 1.3, 1.4, and 1.6.

#### Specific gain

If the heights of channels that are closest to a target stimulus are increased more than distal channels, then the change is referred to as specific gain (see Figure 3). At the target, *r*_*max*_ was multiplied by the the same scaling factors as used for nonspecific gain (see above), but for non-target stimuli the scaling factor was linearly reduced with distance from the target, as described in the introduction to this section.

#### Nonspecific tuning

If the width of the tuning functions is uniformly narrowed across the entire stimulus domain, then the change is referred to as nonspecific tuning (see Figure 3). We tested scaling factors of 0.6, 0.75, and 0.9, which simply multiplied the parameter *ω*_*c*_ for all channels.

#### Specific tuning

If the widths of channels that are closest to a target stimulus are narrowed more than distal channels, then the change is referred to as specific tuning (see Figure 3). At the target, *ω*_*c*_ was scaled by factors of 0.3, 0.4, and 0.6, but for non-target stimuli the scaling factor was linearly reduced with distance from the target, as described in the introduction to this section.

#### Inward tuning shift

If each channel’s preferred stimulus moves toward the target stimulus, then the change is referred to as an inward tuning shift (see Figure 3). The shifting of the parameter *s*_*c*_ for a channel exactly next to the target were −10, −15, and −20 for simulations involving the sparse population, and −5, −7.5, and −10 for simulations involving the dense population, where the “–” sign refers to shift towards the target. The dense populations had reduced parameters to prevent channels from crossing/overshooting the target. With a circular domain, there are no true “nonspecific” tuning shifts, as the movement of channels toward the target must necessarily remove channels in areas away from the target. Thus, we simulated only “specific” versions of tuning shift, in which the magnitude of the shift in *s*_*c*_ was linearly reduced with distance from the target, as described in the introduction to this section.

#### Outward tuning shift

If each channel’s preferred stimulus moves away from the target stimulus, then the change is referred to as an outward tuning shift (see Figure 3). The shifting of the parameter *s*_*c*_ for a channel exactly next to the target were +10, +15, and +20 for simulations involving the sparse population, and +5, +7.5, and +10 for simulations involving the dense population, where the “+” sign refers to shift away from the target. With a circular domain, there are no true “nonspecific” tuning shifts, as the movement of channels away from the target must necessarily concentrate channels in areas away from the target. Thus, we simulated only “specific” versions of tuning shift, in which the magnitude of the shift in *s*_*c*_ was linearly reduced with distance from the target, as described in the introduction to this section.

#### Specific suppression

If the height of tuning functions for channels that are closest to the target stimulus are decreased less than for distal channels, then the change is referred to as specific suppression (see Figure 3). Specific suppression can be thought of as the inverse of specific gain: while the target channel’s responsiveness is unaffected, the maximum responses of the remaining channels are reduced as a function from their distance to the target (with a larger distance leading to greater suppression). At the target, *r*^*max*^ was left the same, but for channels with non-target preferred stimuli the scaling factor was linearly reduced with distance from the target, as described in the introduction to this section. An additional scaling parameter was necessary to determine the maximum level of suppression in *r*^*max*^ (or, conversely, the minimum value toward which *r*^*max*^ was linearly reduced). These scaling parameters were obtained by inverting the corresponding values used for the gain simulations (i.e., ^1^/_1.3_, ^1^/_1.4_, ^1^/_1.6_; see rounded values in Table 3).

#### Specific suppression plus gain

Specific suppression plus gain is the only combination of two mechanisms of encoding change that we implemented in our simulations (see Figure 3). The reason was that it has been specifically proposed in the literature as a way in which selective attention improves stimulus encoding [3]. To implement this combination, we used the highest level of suppression from the previous simulation (i.e., ^1^/_1.6_ = 0.63) and used three different levels of nonspecific gain: 1.4, 1.3, and 1.2.

### Simulated Psychophysical Functions

#### Threshold vs Noise (TvN) curve

In these simulations, we obtained thresholds from each model under study as explained above. At each repetition *m* = 1, 2, …*M*, we presented the model with both the target stimulus and a pattern of external noise with a certain magnitude . We assumed that the pattern of external noise had two effects on the response of the model. First, because a pattern of external noise can sometimes approximate each channel’s preferred stimulus to a certain extent, we stimulated each channel with its preferred stimulus and then scaled down the population response through a random weight. Each weight was randomly sampled from a normal distribution with mean zero and a standard deviation *σ*_*E*_, which was varied to produce different levels of external noise. The second effect of the pattern of external noise would be to degrade the presented target stimulus (e.g., see grating stimuli in Figure 9). Because the target cannot be represented perfectly in the presence of external noise, its produced population response was also scaled by a weight equal to 1 − *σ*_*E*_.

There were 375 levels for the external noise parameter *σ*_*E*_, ranging from 0.0 (no external noise) to approximately 0.5 (0.513 a point where the noise response vectors had approximately the same strength as the target). The levels of external noise were evenly spaced along a log scale, meaning that values were closer together around 0 and because increasingly more separated as they approached the maximum.

After obtaining thresholds at each of the 375 levels of external noise, B-spline smoothing was applied to the resulting TvN curves, in an attempt to obtain better approximations to an “idealized” TvN curve from each model. For all TvN curves, a smoothing parameter of 50 was selected. The goal behind parameter selection was to represent the ideal (i.e., smooth) curve as closely as possible without disrupting the relative threshold relations between the curves.

The dense population models in general produced lower thresholds than the sparse population models. To aid comparison across models, we transformed the TvN curves obtained using sparse models, so that they would be in the scale shown by curves obtained using dense models. We obtained a scaling vector by dividing the sparse baseline TvN by the dense baseline TvN, after smoothing, at each of the 375simulated levels of external noise. This scaling vector was then applied to every sparse TvN curve, by applying the Hadamard product between both vectors.

#### Threshold vs Stimulus (TvS) curve

In these simulations, we presented 181 stimuli, ranging evenly from −90 to 90, to each variation of the model. No external noise was presented, but neural noise was still present. As before, B-spline smoothing was applied to the obtained TvS curves. Because the stimulus domain is circular, the thresholds also were appended to both sides to prevent the B-splines from curling at the ends. We used different smoothing parameters for different simulations, again with the goal of representing the ideal (i.e., smooth) curve as closely as possible without disrupting the relative threshold relations between the curves. For the dense populations, the standard smoothing weight was 3, but nonspecific tuning only needed a weight of 2, and specific suppression needed a weight of 4. For the sparse populations, the standard smoothing weight was 6, but nonspecific tuning only needed a weight of 4, specific suppression needed a weight of 11, inward shift needed a weight of 7, and specific gain needed a weight of 50.

To aid comparison across models, we transformed the TvS curves obtained using sparse models, so that they would be in the scale shown by curves obtained using dense models, in the same way as described for TvN curves above.

### Curve-Fitting Analysis of Population Responses

For homogeneous populations, the population response has the same shape as the common channel tuning function. However, with non-homogeneous populations such as those obtained after applying an encoding change mechanism, it is possible that the population response might not be well-described by the function used to model tuning functions. For this reason, we fitted population responses obtained from the model to a function which has the Gaussian as a special case, but that can also produce curves with lower or higher curvature at the peak than the Gaussian (including exponential functions, when the curvature at the peak is very low). The function is defined by the following equation:

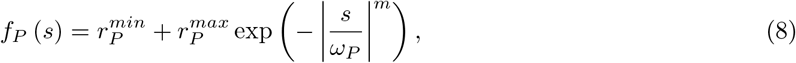
where 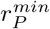 is the baseline level of responding (set to zero, as no observed curves showed higher baseline), 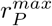 is the peak level of responding, *ω*_*P*_ determines the width of the function (the inverse of its decay slope), *m* determines the width of the function at the peak (with *m* = 2 representing a Gaussian function, and *m* = 1 representing an exponential decay function), and *s* is the stimulus value. A similar function has been fitted to estimates of population responses obtained using inverted encoding modeling [38].

The population curve in Equation 8 was fit to the population responses obtained from the model using least squares estimation with the bound-constrained optimization method (L-BFGS-B) included with *optim* in R v. 3.5.1. The lower bounds enforced for the three fitted parameters were as follows: 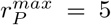, *ω*_*P*_ = 1, and *m* = 1. The same values were provided as starting parameters for the optimization algorithm. Whenever the optimization algorithm did not converge, the optimization was performed again with a new set of starting parameters, equal to the previous starting parameters plus random values obtained from a normal distribution with mean zero and variance equal to five. This procedure was repeated until a satisfactory converging solution to the minimization problem was obtained. The results from this analysis (Figure 7) were plotted using *ggplot2* v. 3.1.

The fitted population response curves were compared against smooth population responses directly obtained from a model with an very dense population of 200 channels, with position parameters distributed homogeneously along the stimulus dimension. We used the squared of the Pearson correlation, *r*^2^, between the two curves to evaluate how well the population curve fitted to sparse data captured features of a fine-grained population response.

### Simulation environment specifications

The simulations were run on a Titan W375 Workstation PC with 32 dual-core (64 cores and 128 threads total) AMD EPYC 7551 2.0GHz (3.0GHz Turbo) 64MB Cache processors, running Ubuntu 18.04.4 LTS. Simulations were programmed using Python 3.7.2, extended with numpy v. 1.15.4, scipy v. 1.1.0. For parallelization, ipyparallel v. 6.2.3 was used in conjunction with jupyter-client v. 5.2.4 and notebook v. 5.7.4 integration. The pandas v. 0.23.4 module was used to save, read, and manage data, and matplotlib v. was used to plot data. Python and all modules were obtained through the Anaconda distribution v. 4.5.12.

